# A JAK-STAT-activated myeloid hub at ulceration sites underlies resistance to anti-TNF therapy in Crohn’s disease

**DOI:** 10.64898/2026.02.12.705345

**Authors:** Thomas Laurent, Monika Mykhaylyshyn, Jake R. Thomas, Gaelle Beriou, Lucas Otero-Laudouar, Lucas Brusselle, Aurelie Joussaume, Cecile Braudeau, Justine Chevreuil, Cynthia Fourgeux, Laurence Delbos, Martin Braud, Nicolas Chapelle, Mathieu Uzzan, Pablo Canales-Herrerias, Theo Soude, Cecile Girard, Jean-François Mosnier, Catherine Le Berre, Juliette Podevin, Emilie Duchalais-Dassonneville, Ángela Sanzo-Machuca, Azucena Salas, Jean-Frederic Colombel, Jeremie Poschmann, Saurabh Mehandru, Arnaud Bourreille, Ephraim Kenigsberg, Miriam Merad, Andreas Schlitzer, Jerome C. Martin

**Affiliations:** Nantes Université, CHU Nantes, Inserm, Center for Research in Transplantation and Translational Immunology, UMR 1064, F-44000, Nantes, France; Biology of Inflammation, Life & Medical Sciences Institute, University of Bonn, Bonn, Germany; CHU Nantes, Nantes Université, Laboratoire d’Immunologie, Centre d’Immunomonitorage Nantes Atlantique, F-44000, Nantes, France; Paris Est Créteil University UPEC, Assistance Publique-Hôpitaux de Paris (AP-HP), Henri Mondor Hospital, Gastroenterology department, Créteil F-94010, France; Precision Immunology Institute, Icahn School of Medicine at Mount Sinai, New York, NY, USA; Henry D. Janowitz Division of Gastroenterology, Department of Medicine, Icahn School of Medicine at Mount Sinai, New York, NY, USA; CHU Nantes, Nantes Université, Hépato-Gastro-Entérologie et Assistance Nutritionnelle, Inserm CIC 1413, Institut des Maladies de l’Appareil Digestif (IMAD), F-44000, Nantes, France; CHU Nantes, Nantes Université, Anatomie et Cytologie Pathologiques, F-44000, Nantes, France; Nantes Université, CHU de Nantes, Inserm, The Enteric Nervous System in Gut and Brain Disorders, IMAD, F-44000, Nantes, France; Nantes Université, CHU Nantes, Chirurgie Digestive, Institut des Maladies de l’Appareil Digestif (IMAD), F-44000, Nantes, France; Inflammatory Bowel Disease Unit, Hospital Clínic of Barcelona, Institut d’Investigacions Biomediques August Pi i Sunyer, Barcelona, Spain; Department of Immunology and Immunotherapy Icahn School of Medicine at Mount Sinai, New York, NY, USA

## Abstract

Despite major advances in immunotherapies, durable disease control in immune-mediated inflammatory diseases (IMIDs) is frequently limited by resistance driven by maladaptive immune programs, resulting in progressive tissue damage. Myeloid cells are central effectors across IMIDs, including inflammatory bowel disease (IBD), yet how discrete myeloid states govern disease progression and therapeutic response remains incompletely defined. Here, using integrative single-cell transcriptomic and molecular analyses of inflamed human ileum in Crohn’s disease (CD), we identify combinatorial transcriptional programs that revealed previously unrecognized axes of monocyte-macrophage (mo-mac) heterogeneity. These myeloid states and programs were conserved and validated across multiple independent cohorts. We delineate a pathogenic mo-mac subset, interferon-associated inflammatory monocytes (IFIM), shaped by convergent NF-κB and IFNγ signaling and enriched in advanced disease. Baseline abundance of IFIM programs predicted resistance to anti-TNF therapy and severe postoperative recurrence, with IFIM-derived ligands sustaining epithelial injury. IFIM exhibited robust JAK–STAT activation, indicating susceptibility to JAK inhibition with clinically approved agents, including upadacitinib. Exemplified by IFIM-driven inflammation, these findings identify pathogenic myeloid state remodeling as a molecular basis of anti-TNF resistance and provide a framework for mechanism-guided therapeutic strategies across chronic inflammatory diseases.

## INTRODUCTION

Inflammatory bowel disease (IBD), which includes Crohn’s disease (CD) and ulcerative colitis (UC), is characterized by intermittent chronic inflammation of the gastrointestinal tract responsible for bowel damage and disabilities^1^. Triggering events in IBD involve the interplay between environmental factors and genetic susceptibilities^2–4^, which culminate in uncontrolled immune responses against luminal gut antigens^2,5–9^. The unprecedented success of anti-TNF antibodies (Abs)^10^ has led to the realization that targeting dysregulated states of intestinal immunity to achieve strict control of mucosal inflammation, was key to prevent the accumulation of bowel damage and disease progression^11,12^. However, despite the expanding arsenal of immune-targeted therapies in IBD, response rates remain suboptimal, leaving a substantial proportion of patients in need of treatment^13^. Recent studies have provided important insights into the cellular and molecular contexture of human IBD intestinal lesions^14–23^, showing that distinct immune and stromal modules underlie primary resistance to anti-TNF Abs^14–18,24^. These findings clearly indicate that the limited efficacy of current immunotherapies partly reflects an incomplete understanding of the molecular underpinnings driving immunopathological heterogeneity. Addressing these knowledge gaps is therefore critical for establishing a molecular framework to guide immunotherapy decisions and optimizing early combination strategies^13,25^.

Macrophages in the human intestinal *lamina propria* arise from infiltrating classical monocytes and are specialized to maintain homeostasis, resolve inflammation, and promote tissue repair^26,27^. When dysregulated, however, these intestinal monocyte-derived macrophages (mo-macs) drive chronic, self-perpetuating inflammation^28–34^. GWAS and risk allele-gene expression studies further implicate macrophage dysfunction in severe IBD. Both macrophage accumulation and phenotypic alterations have been widely reported in CD^3,35–43^, and our prior studies have suggested central roles for macrophages in orchestrating refractory inflammation in CD^15^. Nevertheless, the underlying mechanisms and extrinsic cues altering monocyte-to-macrophage trajectories associated with CD inflammation in the human ileum remain poorly understood^14,15,33,36,37,44^. A major challenge has been the absence of a clear framework defining mo-mac cell states and expression programs, including how these programs shape inflammatory responses and clinical outcomes. Standard single-cell analyses indeed often fail to distinguish maturation-associated programs from functional immune states, limiting resolution of transitional and heterogeneous mo-mac populations.

In this study, we developed a new analytical framework to resolve the molecular heterogeneity of mo-mac states in ileal CD. This approach uncovered a previously unrecognized pathogenic mo-mac state that drives resistance to anti-TNF therapy and underlies treatment-refractory inflammation.

## RESULTS

### Program-based combinatorial definition of mo-mac states in CD ileums

We sought to capture mo-mac programs in severe CD at the highest resolution in a discovery analysis by combining deep profiling of mononuclear phagocytes (MNPs) with a high-dimensional modeling approach (**Fig. 1A**). Purified MNPs were isolated from the inflamed, surgically resected terminal ileum of seven patient included at CHU Nantes and analyzed by single-cell RNA sequencing (scRNA-seq) (**Fig. S1A**; **Tables S1&2**). Using the Metacells pipeline^45,46^, we reconstructed the transcriptional landscape of MNPs in CD, identifying multiple discrete and continuous cellular states that were coherent with canonical gene programs defining known MNP subsets (**Fig. S2A**). Gene module–based analysis^19^ of the metacell manifold revealed an additional layer of combinatorially expressed transcriptional modules, uncovering previously unrecognized axes of myeloid heterogeneity within the inflamed ileum (**Fig. S2B, Table S3, methods**). In total, we curated and annotated 17 gene programs and leveraged their combinatorial expression patterns to define nine mo-mac molecular states with consistent program-derived gene signatures (**Fig. 1B-C, S2C**; **Table S4**). We validated these transcriptional programs and monocyte-macrophage states in an independent scRNA-seq dataset from a larger cohort of patients with advanced disease from CHU Nantes and Mount Sinai Hospital (referred to as the CHU Nantes/Mount Sinai Hospital cohort) (**Fig. S3A&B; Table S5**)^15,19^.

**Figure 1:**
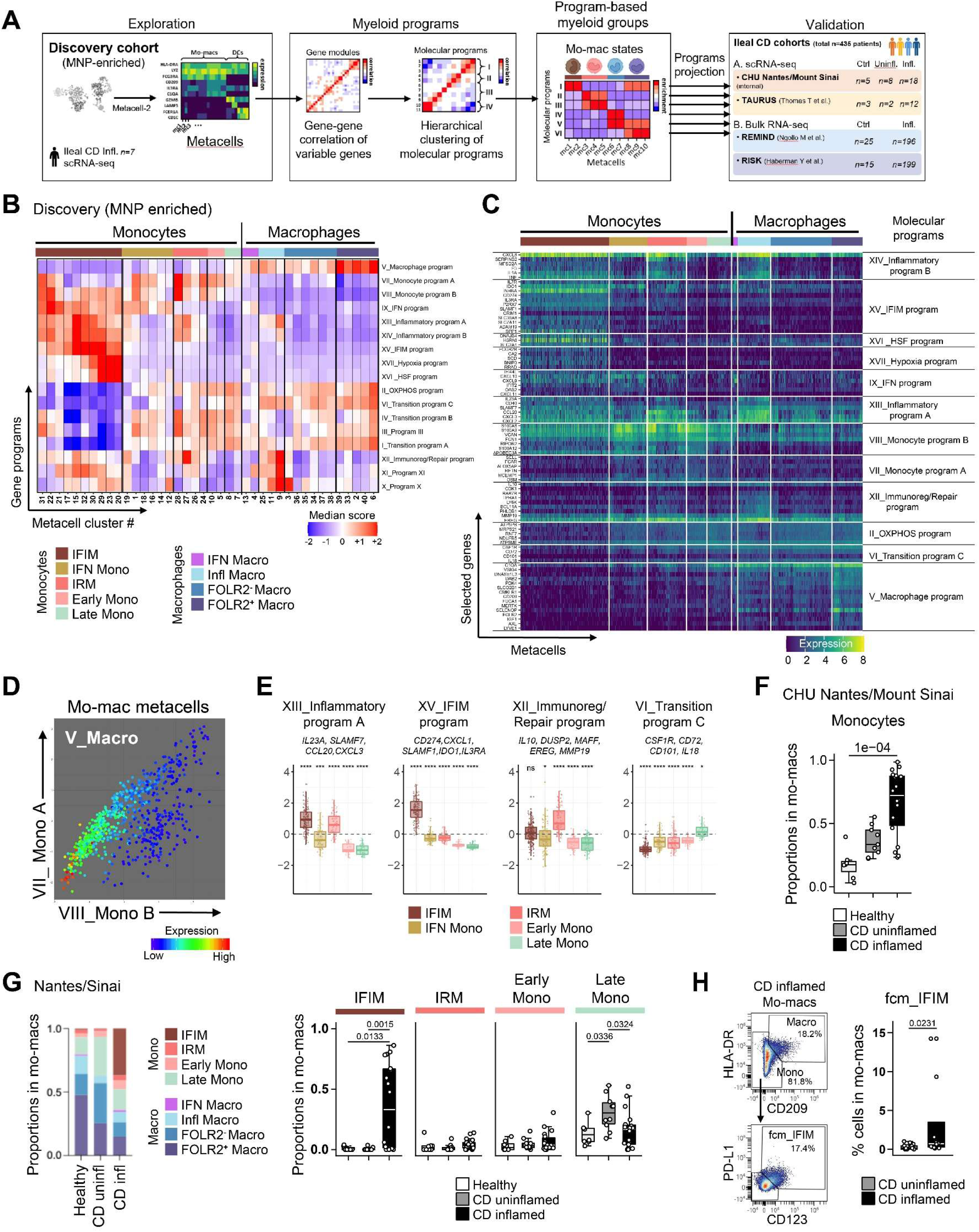
Program-based combinatorial definition of mo-mac states in CD ileums. **(A)** Workflow of the program-based combinatorial analysis of monocytes/macrophages present in the inflamed ileum of patients with severe CD. **(B-D)** MNP were flow-sorted from inflamed ileums of CD patients with stricturing disease (n=7) and analyzed by scRNAseq with Metacells-2. (B) Heatmap showing expression of the 17 molecular mo-mac programs (rows) in metacell clusters (columns) grouped according to combinatorial profiles into 9 molecular states of mo-macs. (C) Heatmap showing expression of representative genes from mo-mac programs (rows) in single metacells (columns) grouped as defined in B. (D) Scatterplot of mo-macs metacells according to their expression of the two monocyte gene programs (Monocyte program A – y-axis; Monocyte program B – x-axis) and the macrophage program (Macrophage program– color-coded). **(E)** Boxplots comparing expression levels of indicated programs across the five monocyte groups. Selected representative genes from the programs are indicated. P-values were calculated using Wilcoxon signed-rank tests (group median vs. global mean, **Table S21**) with Bonferroni correction between all groups. *\*P < 0.05, **P < 0.01*, *\*\*\*P < 0.001, ****P < 0.0001, n.s. non-significant.* **(F)** Boxplots comparing the proportions of monocytes in total mo-macs in the ileum between indicated groups. Each dot represents an ileum sample, and lines represent median and quartiles. The whiskers represent 1.5 times the interquartile range. P-values were obtained by Dunn’s post hoc test following Kruskal–Wallis, adjusted with Holm correction for multiple testing (**Table S20**). **(G)** Bar graphs showing monocyte and macrophage profiles among indicated groups (left). Each bar corresponds to the averaged proportion in each group. Right: Boxplots showing the proportion of monocyte groups in total mo-macs among indicated groups. Each dot represents a single ileum sample, and lines represent median and quartiles. The whiskers represent 1.5 times the interquartile range. P-values were obtained by Dunn’s post hoc test following Kruskal–Wallis, adjusted with Holm correction for multiple testing (**Table S22**). **(H)** Representative FACS plots showing the proportions of monocytes/macrophages in total mo-macs and FCM_IFIM in monocytes from inflamed CD ileums (left). Right: Boxplots showing the proportions of FCM_IFIM in total mo-macs among indicated groups. Each dot represents a single ileum sample, and lines represent median and quartiles. The whiskers represent 1.5 times the interquartile range. P-values were calculated using a Mann-Whitney *U* test (**Table S23**).

We classified the nine mo-mac groups as five monocyte and four macrophage states based on their expression of three general “identity” gene programs: **“Monocyte_program_A”** (prog. VII; *SELL*, *CD55*, *FCAR*, *RETN*), **“Monocyte_program_B”** (prog. VIII; *S100A8/A9*, *S100A12*, *VCAN*, *FCN1*) and **“Macrophage_program”** (prog. V; *C1Qs*, *CD209*, *MERTK*, *FUCA1, SELENOP*). The two monocyte gene programs were frequently, but not systematically, co-expressed and gradually decreased as the macrophage program was induced (**Fig. 1D, S3C**). In the CHU Nantes/Mount Sinai cohort, which included inflamed (n=20), non-inflamed (n=10) and healthy (n=8) ileal samples analyzed by scRNA-seq, monocytes were enriched in inflamed tissues, comprising approximately two-thirds of the mo-mac compartment (**Fig. 1F**). This marked predominance positioned monocytes as the principal component shaping the mo-mac compartment in severe CD.

Reflecting their high plasticity^47^, the molecular diversity of monocytes in the CD-inflamed ileum could not be captured by a single linear trajectory (**Fig. S3D**). Accordingly, we identified two transcriptionally divergent inflammatory states, despite both sharing the same inflammatory programs: **“Inflammatory_program_A”** (prog. XIII *IL23A*, *IL6*, *CXCL2*, *CXCL3*) and **“Inflammatory_program_B”** (prog. XIV *IL1B*, *TNF*, *CCL3*, *CXCL8*) (**Fig. 1E, S2C**). The first state was annotated as interferon-associated inflammatory monocytes (IFIM) based on co-expression of an interferon signature (**“Interferon”** prog. IX; *IRF1*, *CXCL10*, *IFIH1*, *GPB4*, *WARS1*, *CIITA*, *OAS2*). In addition, IFIM uniquely expressed a distinct transcriptional module, the **“IFIM-program”** (prog. XV), containing genes involved in neutrophil production and recruitment (*CXCL1*, *CXCL5*, *CSF3*)^48^, immunoregulation (*IDO1*, *CD274*)^49^, and broader pro-inflammatory functions (*CCL5*, *INHBA*, *SLAMF1, IL3RA*)^50–52^ (**Fig. 1E, S2C**). IFIM constituted the predominant monocyte population in the CHU Nantes/Mount Sinai cohort (**Fig. 1G**). By contrast, the second inflammatory monocyte state was annotated as immunoregulation-repair monocytes (IRM), as it induced program **“Immunoregulation/Repair”** (prog. XII; *IL10*, *NFKBID, OLR, EREG*, *MMP19*) but lacked the **“IFIM-program”**. Accordingly, unlike IFIM, IRM represented only a minor group in the inflamed ileum of severe patients (**Fig. 1G**). Beyond IFIM and IRM, remaining monocyte groups included IFN_Mono, which were rarely detected in the CHU Nantes/Mount Sinai cohort, as well as early and late maturation-stage monocytes distinguished by differential expression of **“Transition”** programs and lacking inflammatory programs (**Fig. 1E, 1G**).

To validate the accumulation of the IFIM monocyte state in the CHU Nantes/Mount Sinai cohort of severe CD patients, we used *CD274* (PD-L1) and *IL3RA* (CD123), two membrane markers from the **“IFIM-program”** (**Fig. S4A-B**). Cytometry analyses confirmed the enrichment of PD-L1^+^ CD123^+^ mo-macs in inflamed ileum from CD patients analyzed at Mount Sinai Hospital by mass cytometry (CyTOF^15^) and CHU Nantes by spectral flow cytometry (spectral FCM). In the latter, detailed phenotypic characterization further confirmed PD-L1^+^ CD123^+^ mo-macs as CD209^-^HLA-DR^low^ monocytes (referred to as FCM_IFIM to mirror scRNA-seq annotations) (**Fig. 1H, S1B-C, S4C-E, Table S6-9**). Finally, acknowledging that mo–mac groups represent continuous rather than discrete states, we leveraged scRNA-seq expression of *HLA-DRA*, *ITGAM*, *FOLR2* and *CD163* (**Fig. S2D**) to define gating strategies for FCM_mo-mac populations and broaden their phenotypic characterization in CD (**Fig. S1C, S4E-F, Tables S7-9**).

Altogether, our program-based molecular dissection reveals a granular and previously underappreciated diversity of the mo–mac compartment in the CD-inflamed ileum and identifies IFIM as a distinct inflammatory monocyte state that dominates this compartment in severe disease.

### Enrichment in IFIM associates with CD severity

In CD, molecular heterogeneity remains a major barrier to implementing mechanism-guided immunotherapy early in the disease course^15^. Although IFIM enrichment was prominent in most patients with severe disease, it remained heterogeneous across individuals (**Fig. 2A**). To determine whether IFIM represents a myeloid response linked to therapeutic resistance and disease progression, we therefore integrated three additional publicly available single-cell and bulk transcriptomic datasets respectively from the TAURUS^23^, REMIND^53^ and RISK^55^ cohorts (**Fig. 1A**).

**Figure 2:**
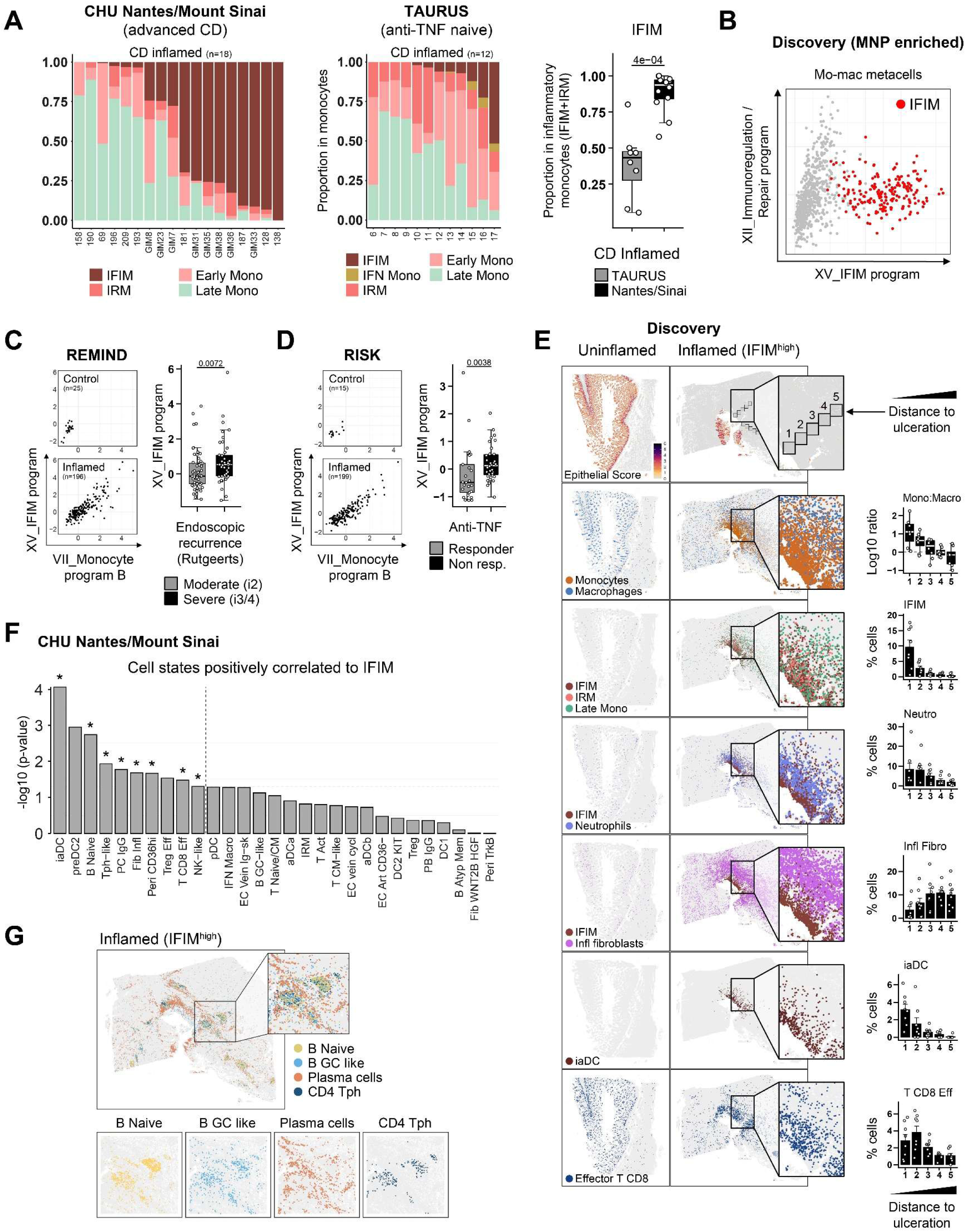
Enrichment in inflammatory IFIM at sites of severe epithelial damage. **(A)** Bar graphs showing monocyte profiles across patients in the Mount Sinai/CHU Nantes cohort (left) and in the TAURUS cohort (middle). Each bar corresponds to one patient. Right: Boxplots comparing the proportion of inflammatory IFIM in total inflammatory monocytes (IFIM+IRM) between CD patients in both cohorts. Each dot represents a CD patient, and lines represent median and quartiles. The whiskers represent 1.5 times the interquartile range. P-values were calculated using a Mann-Whitney *U* test (**Table S24**). **(B)** Scatterplot of total mo-mac metacells according to their expression of programs “Immunoregulation/Repair” (y-axis) and “IFIM-program” (x-axis). Red dots correspond to IFIM metacells. **(C)** Scatterplots showing expressions of programs “Monocyte program B” (x-axis) and “IFIM-program” (y-axis) in ileums profiled in bulk from CD patients in the REMIND cohort (n=221 total; ctrl n=25; CD=196) (left). Right: Boxplots comparing baseline expression of program “IFIM-program” in patients who experienced clinical recurrence 6 months post-resection. Each dot represents a CD patient. P-values were calculated using a Mann-Whitney *U* test (**Table S25**). **(D)** Scatterplots showing expressions of programs “Monocyte program B” (x-axis) and “IFIM-program” (y-axis) in ileums profiled in bulk from CD patients in the RISK cohort (n=214 total; uninfl. n=15; infl. n=199) (left). Right: Boxplots comparing baseline expression of program “IFIM-program” in newly diagnosed patients treated with anti-TNF (right). Each dot represents a CD patient. P-values were calculated using a Mann-Whitney *U* test (**Table S25**). **(E)** Spatial visualization of epithelial, immune and stromal cell localization in uninflamed (left) and inflamed (middle) ileum cross-sections. Boxplots comparing cell proportions across five sequential bins starting from the ulceration region (bin 1) and progressing deeper in the *lamina propria* in IFIM^high^ patients (n=2 patients, 2 sections per patient) (right). Each dot represents the cell proportions calculated in 500 cells randomly sampled per bin, and lines represent median and quartiles. The whiskers represent 1.5 times the interquartile range. Top to bottom: Quantification of the epithelium gene score, and sequential bins; Monocytes and macrophages, and per-bin quantification of the monocyte/macrophage log10 ratio; Indicated monocyte states and per bin quantification of IFIM; Spatial visualization and per bin quantification of neutrophils, inflammatory fibroblasts, inflammatory activated dendritic cells (iaDC) and effector CD8 T cells. **(F)** Bar plots showing scRNA-seq-defined cell states positively correlated with IFIM in inflamed ileal tissue from CD patients. Cell states above the dashed line have a P-value < 0.05. * denote cell states significantly correlated with IFIM but not with IRM (**Table S26**). **(G)** Spatial visualization and zoomed-in localization of TLS structures including naïve B cells, germinal center (GC)-like B cells, plasma cells and T CD4 peripheral helper (Tph) cells.

We first analyzed data from the IBD TAURUS cohort (Oxford, UK), which enrolled biologic-naïve adult patients with IBD who were escalated to anti-TNF antibody therapy and followed longitudinally, with intestinal biopsies collected for scRNA-seq profiling^23^. Applying our program-based combinatorial analysis, we again recapitulated the nine mo-mac states identified in the discovery analysis and the CHU Nantes/Mount Sinai Cohort (**Fig. S5A-C**). As observed in advanced CD, monocytes already dominated the mo-mac compartment in inflamed ileum samples (**Fig. S5D**). Remarkably, IFIM enrichment was detected across inflamed ileal samples obtained prior to anti-TNF therapy, albeit with substantial interpatient heterogeneity, indicating that IFIM represent an inducible inflammatory monocyte state that can emerge in a subset of patients already at early stages of CD (**Fig. 2A, S5E**). Although limited by sample size, patients who subsequently failed to respond to anti-TNF therapy showed higher IFIM enrichment (**Fig. S5F**). Consistent with an association between baseline IFIM abundance, anti-TNF resistance and progression to more severe disease, the IFIM/IRM ratio was markedly increased in anti-TNF non-responders in the CHU Nantes/Mount Sinai cohort compared with anti-TNF-naïve patients from the TAURUS cohort (**Fig. 2A**). Importantly, the “**IFIM-program”** and **“Immunoregulation/Repair”** molecular programs were consistently expressed in a mutually exclusive manner across mo–mac metacells, confirming IFIM and IRM as two non-overlapping states of inflammatory monocytes in the CD-inflamed ileum (**Fig. 2B, S5G**).

Next, we evaluated the association between IFIM enrichment and CD severity using two publicly available bulk transcriptomic datasets of mucosal ileum. The REMIND cohort prospectively enrolled adult patients with ileal or ileocolonic CD undergoing their first CD-related surgical resection and followed them longitudinally to assess endoscopic recurrence^53^. We assessed IFIM enrichment in resected ileum samples by projecting microarray data onto two combined molecular programs defining the IFIM state (**“Monocyte_program_B”** and **“IFIM-program”**). As expected, expression of both programs was increased in inflamed versus uninflamed ileum and correlated positively (**Fig. 2C**). Among patients who experienced clinical recurrence 6 months post-surgery (Rutgeerts score ≥ i2^54^), baseline co-expression of the two programs identified individuals enriched for severe recurrence (Rutgeerts scores i3 and i4; X^2^ P=0.0246) (**Fig. S5H**). Strikingly, expression of the “**IFIM-program**” alone showed the most significant association with future severe clinical recurrence (**Fig. 2C, S5I**). We applied a similar strategy to the RISK cohort, a prospective inception study of pediatric patients with newly diagnosed CD^55^. Among patients who received anti-TNF Ab therapy within the first year after diagnosis, IFIM enrichment at baseline was associated with future non-response to treatment, defined by failure to achieve durable corticosteroid (CS)-free clinical remission 18–24 months post-diagnosis (**Fig. 2D, S5H-I**).

Taken together, these findings demonstrate that the IFIM monocyte state is induced as part of an inflammatory organization enriched in the ileum of CD patients who do not respond to anti-TNF therapy, marking advanced disease progression and heightened risk of severe surgical recurrence.

### IFIM accumulate at sites of severe epithelial damage

The association between IFIM enrichment and CD progression to severity suggested anti-TNF resistance may reside within IFIM-associated programs, prompting further investigation of their role in inflamed ileal tissues. We therefore selected two IFIM^high^ patients of the CHU Nantes/Mount Sinai cohort based on our scRNA-seq data and analyzed their tissue sections using probe-based spatial transcriptomics. For comparison, we also included a patient with lower IFIM enrichment (**Fig. S6A**). In agreement with our results in dissociated cells, macrophages dominated the mo-mac compartment in the *lamina propria* of uninflamed ileum, and followed a spatial pattern aligned with known distributions in the normal ileum^27,33^. Monocytes were also present, but we confirmed their substantial accumulation in inflamed ileum. Notably, in IFIM^high^ patients, this accumulation was especially pronounced in areas adjacent to the lumen, characterized by severe epithelial damage (**Fig. 2E, S6B**). These regions exhibited a marked enrichment of neutrophils, which extended throughout a broader mucosal area from the site of ulceration (**Fig. 2E and S6C**). We next annotated monocyte profiles using our scRNAseq-derived molecular programs. IFIM accumulation was strikingly confined to these ulceration sites. In these regions, IFIM largely outnumbered IRM and Late_Mono, while their relative abundance decreased progressively in deeper mucosal areas (**Fig. 2E and S6D&E**).

Using detailed cell-lineage annotations in the CHU Nantes/Mount Sinai scRNA-seq dataset (**Fig. S7**), we identified additional cell sates that were positively correlated with IFIM, but not with IRM, in inflamed ileum samples (**Fig. 2F, S9A**). These included inflammatory fibroblasts (r=0.54, p=0.027), consistent with prior reports describing their co-localization with neutrophils in ulcerated areas of anti-TNF resistant UC patients^17^. This analysis further revealed significant associations with effector CD8 T cells (r=0.49, p=0.032), NK-like cells (r=0.456, p=0.0495) and a population of recently described inflammatory activated dendritic cells enriched in chronic inflammation across IMIDs (iaDC r=0.793, p=8.61^^-5^)^56^. In the spatial transcriptomic data, we verified the enrichment of effector CD8 T cells, and iaDC in IFIM-associated regions (**Fig. 2E, S6F**). While we also identified cell subsets reminiscent of tertiary lymphoid structures (TLS), including naïve B cells (r=0.667, p=0.00181), IgG plasma cells (r=0.541, p=0.0168) and CD4 T peripheral helper (TPH)-like (r=0.565, p=0.0116), that correlated with IFIM enrichment, the spatial distribution of TLS in IFIM^high^ inflamed ileum did not indicate preferential localization near ulceration sites (**Fig. 2G, S6G**).

In summary, we show that the IFIM monocyte state is induced within a pathogenic cellular hub located at sites of severe epithelial damage in the inflamed ileum of CD patients, comprising neutrophils, inflammatory fibroblasts, effector CD8 T cells, NK cells, and iaDC.

### IFNγ signaling critically shapes inflammatory monocytes toward the IFIM state

To identify molecular signals preferentially driving IFIM over IRM and potentially contributing to anti-TNF resistance at ulceration sites, we leveraged our program-based analysis to infer the activity of candidate transcription factors (TFs) regulating program induction. First, we performed a motif analysis to identify TF binding sites enriched at transcription start sites of genes within each program (**Tables S10-11; methods**). The TF motif with the highest NES scores in inflammatory programs shared between IFIM and IRM (**“Inflammatory_program_A”** and **“Inflammatory_program_B”**) was REL/Nuclear factor kappa-light-chain-enhancer of activated B cells (NF-κB) (**Fig. 3A, S8A**). In program **“Interferon”**, which was induced in IFIM but not in IRM, we identified the expected motifs for the interferon-stimulated response element (ISRE) and signal transducer and activator of transcription (STAT). These findings, combined with the higher *IRF1* mRNA levels in IFIM (**Fig. S8B**) supported a role for the cytokine IFNγ. IFNγ is the most potent inducer of STAT1-dependent *IRF1* mRNA expression^57–59^ and was previously proposed as a driver of inflammatory mo-macs in CD^35^. Consistently, program **“Interferon”** also contained *CIITA*, *CASP5* and *APOL6* (**Table S4**), which are genes preferentially induced by IFNγ in human cells^60^. Together, these observations suggested that combined NF-kB and IFNγ signaling was likely necessary for establishing the IFIM inflammatory state. In strong support, both REL/NF-κB and ISRE were similarly predicted to regulate the **“IFIM-program”** (**Fig. 3A**). We next cross-referenced the TF motifs identified in this analysis with regulons predicted for mo-macs groups using the SCENIC analytical tool, which estimates TF activity based on gene expression^61^ (**Table S12**). This corroborated shared enrichments of *NFKB1*, *RELA* and *RELB* activities in both IFIM and IRM while *STAT1* and *IRF1* regulons were specific to IFIM (**Fig. 3B, S8C**).

**Figure 3:**
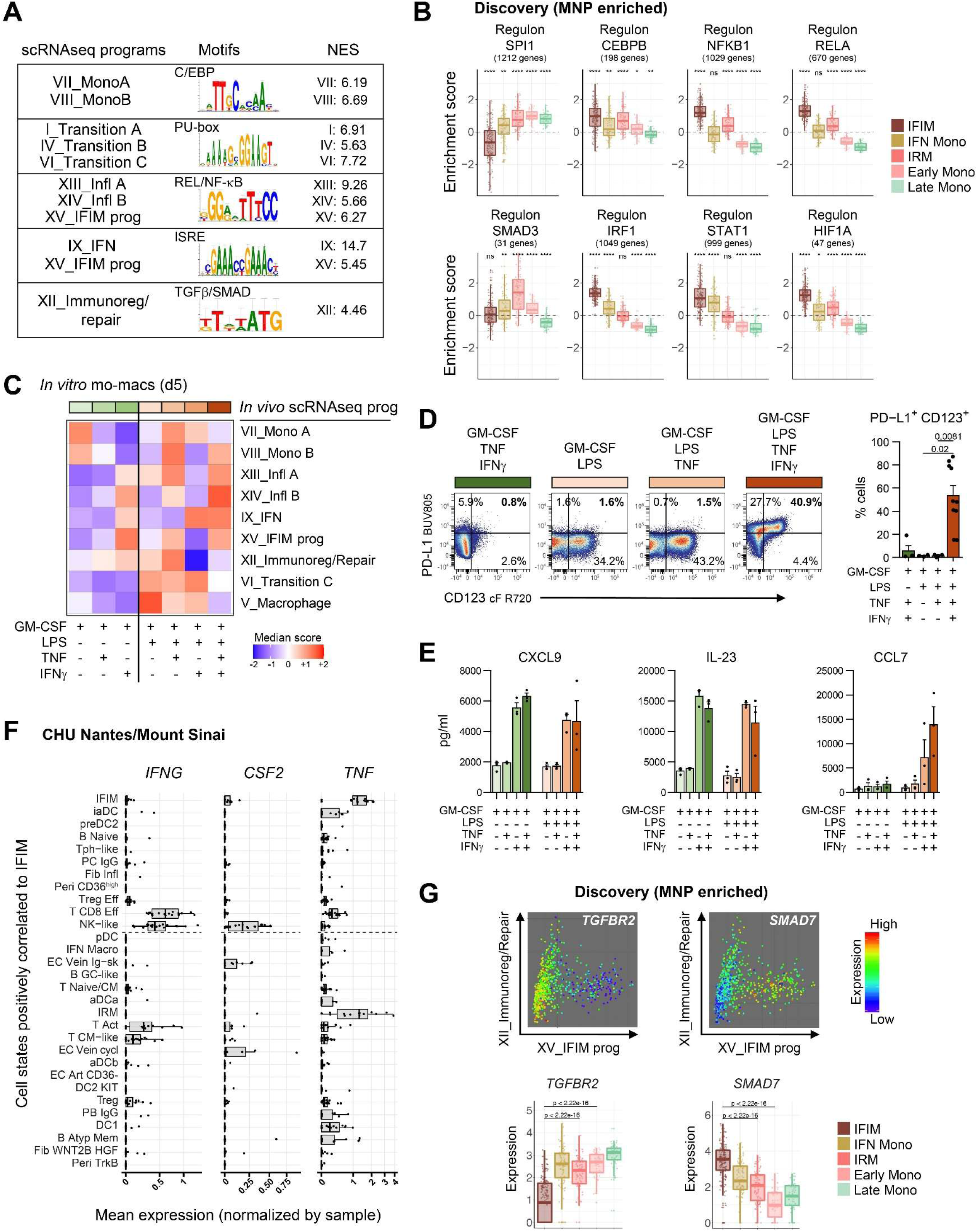
IFNγ signaling shapes the NF-κB-dependent response of inflammatory monocytes toward the IFIM state. **(A)** Top predicted transcription factor (TF) motifs and Normalized Enrichment Score (NES) in the indicated mo-mac programs. **(B)** Boxplots comparing regulon activities of indicated TF calculated by SCENIC between mo-mac groups. P-values were calculated using Wilcoxon signed-rank tests (group median vs. global mean) with Bonferroni correction between all groups. *\*P < 0.05, **P < 0.01*, *\*\*\*P < 0.001, ****P < 0.0001, n.s. non-significant* (**Table S28**). **(C)** Heatmap showing the expression of scRNA-seq-based programs defined *in vivo* in ileal CD mo-macs in blood monocytes from healthy donors analyzed by RNAseq after 5 days of culture *in vitro* in the indicated conditions (n=3). **(D)** FCM scatterplots of PD-L1 and CD123 expressions (left) and bar plots (mean +/− s.e.m.) showing the proportions of PD-L1+ CD123+ cells (right) in monocytes cultured 5 days in the indicated conditions. P-values were calculated using Dunn’s post-hoc pairwise comparisons, with Holm’s correction (**Table S29**). **(E)** Bar plots (mean +/− s.e.m.) showing the concentrations of the indicated cytokines in supernatants collected after 5 days of monocyte culture in the indicated conditions. **(F)** Expression levels of the indicated cytokines across cell states positively correlated with IFIM in inflamed CD ileums. **(G)** Color-coded expressions of *TGFBR2* and *SMAD7* in mo-mac metacells spread onto scatterplots according to their expression of programs “Immunoregulation/Repair” (y-axis) and “IFIM-program” (x-axis) (top). Boxplots comparing expression levels of indicated genes across monocyte groups (bottom). P-values were calculated using Mann-Whitney *U* tests with Bonferroni correction. For clarity, not all comparisons are shown.

To experimentally validate our scRNA-seq-based predictions of the key molecular signals regulating inflammatory IFIM in severe CD *in vivo*, we employed a reductionist *in vitro* system in which highly purified blood monocytes from healthy donors were exposed to ligands selectively activating the predicted pathways. Monocytes were exposed to combinations of IFNγ, lipopolysaccharide (LPS) and TNF for 5 days in the presence of GM-CSF. Using RNA sequencing, we compared the combinatorial expression of the *in vivo* scRNA-seq-based mo-mac programs that best reflected the IFIM state (**Fig. 3C**). While monocytes exposed to GM-CSF alone (mono^GM-CSF(0–120h)^) were primarily defined by co-expression of the two monocyte programs only, combined signaling activities of LPS, TNF and IFNγ best reproduced the IFIM molecular profile. Compared to other conditions tested, Mono^GM-CSF/LPS/IFNγ/TNF(0-120h)^ combined the expression of all three inflammatory programs, including program **“IFIM-program”**, the interferon signature and the monocyte “identity” program **“Monocyte_program_B”** but not **“Monocyte_program_A”**, as well as a lack of induction of **“Immunoregulation/Repair”** or **“Transition”** programs (**Fig. 3C**). Phenotypically, this condition induced coexpression of the IFIM membrane markers PD-L1 and CD123 along with a cytokine secretory profile consistent with scRNA-seq-based predictions for *in vivo* IFIM (**Fig. 3D-E, S1D, S8D-E, Table S13-15**).

Given the central role of IFNγ in inducing the IFIM monocyte state, we next sought to identify its cellular sources at ulceration sites. Analysis of ligand-encoding gene expression across IFIM-associated cell states revealed high levels of *IFNG* in both effector CD8 T cells and rarer NK-like cells. These NK-like cells, along with IFIM, albeit at lower levels, also expressed *CSF2*, which encodes GM-CSF, a major driver of inflammatory mo-mac in human IBD^62,63^, while IFIM and iaDC shared high *TNF* expression with effector CD8 T cells (**Fig. 3F**). Together, these data delineate an IFNγ-dominated cytokine milieu that directs inflammatory monocyte trajectories toward the IFIM state at ulceration sites.

We next asked whether additional signals could also positively promote the IRM state and therefore focused on program **“Immunoregulation/Repair”**. In this program, motif analysis revealed enrichment of motifs associated with TGFβ superfamily signaling (**Fig. 3A, S8A**). Consistently, in IRM, we observed increased regulon activity and mRNA levels of *SMAD3*, a key TF downstream of TGFβs, GDFs, and Activins signaling^64^ (**Fig. 3B, S8B**). Among the genes encoding transmembrane type I and type II receptors for TGFβ ligands, only *TGFBR1* and *TGFBR2* (encoding for TGF-β type I and type II receptors) showed detectable expression levels in mo-macs (**Fig. S8B**). This suggested that *SMAD3* activity in IRM was induced in response to TGFβ signaling. Strikingly, IFIM exhibited markedly reduced mRNA levels of *TGFBR2*, which is critical for initiating receptor-regulated SMAD3-dependent TGFβ signaling in target cells^64^. IFIM, however, upregulated *SMAD7*, a known negative regulator of this pathway (**Fig. 3G, S8B**). While *SMAD7* expression is inducible by TGFβ as part of a negative feedback regulation, it is also upregulated by antagonistic crosstalk with JAK-STAT and NF-kB pathways triggered by cytokines such as IFNγ, IL-6, TNF and IL-1β^65–68^. Concordantly, high expression levels of program “IFIM-program” in mo-macs associated with decreased *TGFBR2* and increased *SMAD7* mRNA expressions, as also verified in both CHU Nantes/Mount Sinai and TAURUS datasets (**Fig. 3G, S8F**). Consistent with our *in vivo* findings, mono^GM-CSF/LPS/IFNγ/TNF(0-120h)^ exhibited the greatest reduction in *TGFBR2* expression compared to mono^GM-CSF(0-120h)^ (**Fig. S8G**).

Finally, corroborating our program annotations based on gene composition, we identified the highest NES scores for CCAAT-enhancer-binding protein (C/EBP) in the two monocyte programs, and for PU-box in the three **“Transition”** programs (**Fig. 3A, S8A**). CEBPB and SPI1 are well-established TFs that respectively bind C/EBP and PU-box to control monocyte identity and monocyte-to-macrophage differentiation^69^. This was supported by higher regulon activity of *CEBPB* in the monocyte groups compared to macrophages (**Fig. 3B**). Consistent with the lack of mo-mac **“Transition”** programs in IFIM, *SPI1* regulon usage was limited to the other monocyte groups. In fact, expressions of programs **“IFIM-program”** and **“Transition”** were anti-correlated across mo-mac metacells (spearman ρ=0.75; P<10^^-10^) (**Fig. S8H**). Additional motifs identified included ETS-like, RAR/RXR and RUNX/AML for the monocyte programs, as wells as ETS, ISRE, RUNX/AML, REL/NF-κB and EGR-site for the “Transition” programs, all proposed to orchestrate monocyte-to-macrophage differentiation (**Fig. S8A, Tables S10-11**)^70^.

Collectively, our results identify a major role for IFNγ signaling provided by CD8 T cells and NK cells in shaping the NF-κB-dependent inflammatory monocyte response toward the IFIM state at sites of severe epithelial damage in the CD ileum.

### An IFIM-centered network sustains a pro-inflammatory hub at ulceration sites

Next, we investigated the contribution of IFIM to the inflammatory cellular hub at ulceration sites by examining their production of specific ligands. Direct comparisons revealed both shared and unique predicted secretomes between inflammatory IFIM and IRM (**Fig. 4A, Table S16**). Notably, IFIM selectively expressed the IFNγ-inducible chemokines *CXCL10*, *CXCL9*, and *CXCL11*. Within the cellular states defining the inflammatory hub, expression of these chemokines was largely restricted to IFIM and iaDC, whereas their receptor *CXCR3* was detected on effector CD8 T cells, NK cells, and IgG plasma cells (**Fig. 4B, S9B**). Consistently, *CXCL9* and *CXCL10* expression was spatially limited to ulceration sites (**Fig. 4C**). Unbiased ligand-receptor analysis further confirmed that these chemokines in monocytes and *CXCR3* in T cells formed an interaction network specific to IFIM-enriched patients (**Fig. S9C**).

**Figure 4:**
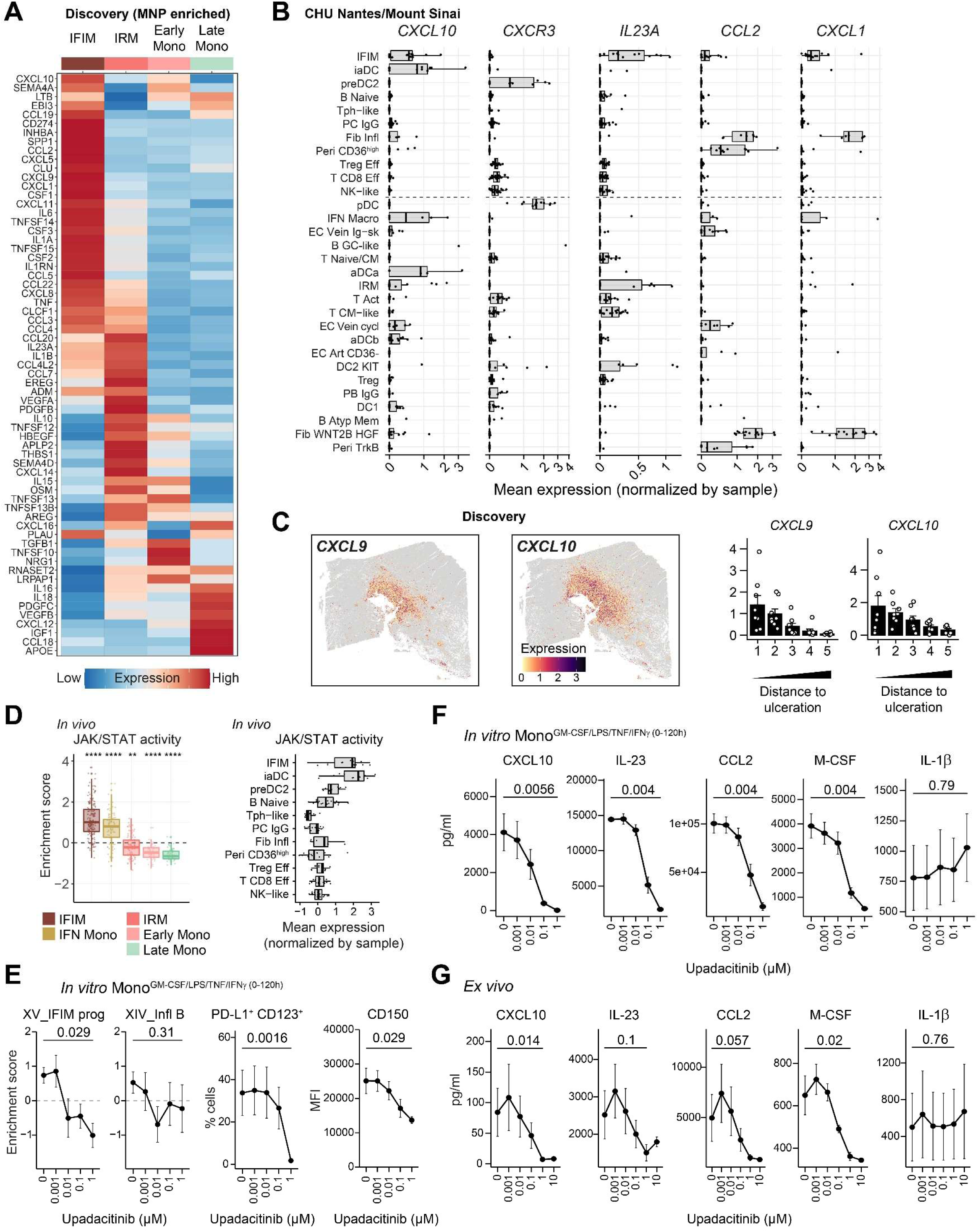
High JAK-STAT activation makes IFIM targetable by JAK inhibitors. **(A)** Heatmap showing the mean expression of indicated genes across monocytes groups **(B)** Expression levels of the indicated genes across cell states positively correlated with IFIM in inflamed CD ileums. **(C)** Spatial distribution of *CXCL9* and *CXCL10* gene expressions in inflamed ileum of IFIM^high^ patients. Boxplots depicting per bin mean expression of the indicated gene starting from the ulceration region as in Fig. 2. **(D)** Left: Boxplots comparing expression levels of predicted JAK/STAT activity across mo-macs groups *in vivo*. P-values were calculated using Wilcoxon signed-rank tests (group median vs. global mean) with Bonferroni correction between all groups. *\*P < 0.05, **P < 0.01*, *\*\*\*P < 0.001, ****P < 0.0001, n.s. non-significant* (**Table S30**). Right: Expression levels of the predicted JAK/STAT activity across cell states positively correlated with IFIM in inflamed CD ileums. **(E)** Monocytes from healthy donors were exposed 5 days to GM-CSF/LPS/TNF/IFNγ and upadacitinib at the indicated concentrations. Dose effect of upadacitinib on the expressions of indicated scRNA-seq-defined mo-mac programs assessed by RNAseq (left), on monocytes phenotypes (middle), and MFI levels of CD150 (right). **(F)** Dose effect of upadacitinib on cytokines levels detected in supernatants of monocytes from healthy donors were exposed 5 days to GM-CSF/LPS/TNF/IFNγ. **(G)** Dose effect of upadacitinib on cytokines levels detected in *lamina propria* cells isolated from surgically resected inflamed ileums of anti-TNF resistant CD patients after 2 days of culture.

Other notable interaction pairs between IFIM and T cell involved *CD274*, *CCL5* and *IL23A*, (encoding for the p19 chain of the cytokine IL-23), a well-established driver of IFNγ in CD8 T cells^60^ (**Fig. 4B, S9C**). Compared with IRM, IFIM also exhibited higher expressions of *CSF1*, a key growth factor for monocyte survival and differentiation in the gut^33^, hence suggesting potential self-amplifying loops (**Fig. 4A)**. *CCL2*, a major monocyte-attractant, was detected in IFIM and at even higher levels in the IFIM-associated stromal populations (**Fig. 4B**). While both IFIM and IRM expressed *CXCL8*, a potent neutrophil-chemoattractant, IFIM uniquely expressed additional genes promoting CXCR2-depdendent neutrophil recruitment (*CXCL1* and *CXCL5*), and neutrophil survival/proliferation (*CSF3* encoding G-CSF), indicating enhanced capacity to drive neutrophil accumulation (**Fig. 4A**). These ligands were also expressed by infl. fibroblasts, suggesting a cooperative role in recruiting monocytes and neutrophils, with IFIM potentially orchestrating their precise positioning at ulceration sites. (**Fig. 4B, S9B**). In contrast to IFIM, the IRM predicted secretome was enriched for ligands associated with immunoregulation and tissue remodeling. Monocyte interactions specific to CD patients lacking IFIM included *IL10* with multiple lineages*, THBS1* with endothelial cells and *HBEGF* with fibroblasts^71,72^ (**Fig. 4A, S9C**). Shared monocyte interactions across both patient groups involved *IL1B*, *CCL18*, *IL15* and *CCL20* with T cells, myeloid and stromal populations.

Together, these data indicate unique functions of IFIM in orchestrating an interaction network supporting activity and maintenance of a complex inflammatory cellular hub at sites of severe ulcerations.

### High JAK-STAT activation makes IFIM targetable by JAK inhibitors

Our results identify IFIM inflammatory programs as central drivers of ulceration-site aggregates in CD that are unlikely to be fully controlled by anti-TNF therapy alone, highlighting them as a potential target for intervention. To pinpoint therapeutically actionable pathways in IFIM, we analyzed pathway activities in mo-mac groups using PROGENy, a method that leverages publicly available perturbation experiments to define pathway-responsive gene signatures^73^. While both IFIM and IRM exhibited elevated NF-κB activity, IFIM uniquely displayed high JAK/STAT activity (**Fig. 4D, S10A-B**). This finding is consistent with our prediction that IFIM are strongly dependent on IFNγ signaling, which activates JAK1 and JAK2 kinases and downstream STAT1^74^. Concordant results were obtained in our *in vitro* system, where mono^GM-CSF/LPS/IFNγ/TNF(0-120h)^ showed the highest JAK/STAT pathway activity compared with other tested conditions, including mono^GM-CSF/IFNγ(0-120h)^ (**Fig. S10C**). Importantly, among IFIM-associated cell states at ulceration sites, iaDC also exhibited elevated JAK/STAT activity, suggesting that therapeutic blockade of this pathway could preferentially target MNP-driven pathogenic functions within the ulceration-associated inflammatory hub (**Fig. 4D**).

Upadacitinib, a JAK inhibitor with high selectivity for JAK1, was recently approved for the treatment of CD patients who have failed anti-TNF therapy^75^. Using our *in vitro* system to assess the ability of upadacitinib to interfere with the induction of the IFIM-like state, we observed not only the expected dose-dependent suppression of program **“Interferon”**, but more importantly a marked inhibition of the **“IFIM-program”** (**Fig. 4E, S10D**). In contrast, the expression of **“Inflammatory_program_B”** was minimally affected, consistent with its lack of direct dependency on IFNγ signaling (**Fig. 4E**). Upadacitinib dose-dependently suppressed the induction of monocytes recapitulating the IFIM phenotype, defined by co-expression of PD-L1, CD123 and CD150, the latter encoded by *SLAMF1* within the **“IFIM-program”** and recently proposed as a marker of inflammatory monocytes in severe fibro-stenotic ileal CD^50^ (**Fig. 4E, S10E; Table S17**).

Guided by our L:R analyses, we next assessed whether pharmacological inhibition of IFIM activity could attenuate the cytokine signals predicted to sustain inflammatory aggregates at ulceration sites. *In vitro*, upadacitinib markedly suppressed CXCL10 secretion by mono^GM-CSF/LPS/IFNγ/TNF(0-120h)^ (**Fig. 4F**). A comparable dose-dependent inhibitory effect was *observed ex vivo* following treatment of lamina propria cells isolated from inflamed ileum of anti-TNF–refractory CD patients (**Fig. 4G**). Beyond CXCL10, upadacitinib consistently reduced secretion of the majority of ligands predicted to mediate IFIM-specific functions, both *in vitro* and *ex vivo* (**Fig. 4F-G, S10F-G; Tables S18-19**). These included IL-23, previously implicated in mo-mac-dependent IFNγ production by T lymphocytes in CD^35^, as well as CCL2, CCL7 and CSF1, predicted to support self-amplifying accumulation of inflammatory monocyte at ulceration sites (**Fig. 4F-G, S10F-G; Tables S18-19**). By contrast, secretion of cytokines not directly linked to IFNγ-dependent transcriptional programs in monocytes, including IL-1α, IL-1β and IL-6, was largely unaffected by upadacitinib **(Fig. 4F-G, S10F-G; Tables S18-19**), indicating selective suppression of IFIM-associated inflammatory outputs.

Collectively, these results define an IFIM-driven molecular mechanism orchestrating inflammatory cellular hubs at sites of severe ulceration in ileal CD, which resist anti-TNF therapy alone and are selectively targetable with upadacitinib.

## DISCUSSION

Molecular heterogeneity is increasingly recognized as a major determinant of immunotherapy success^76^. This underscores the need for improved patient stratification and the development of molecularly guided therapies to optimize outcomes in early-stage CD. Although immune and stromal heterogeneity have been linked to primary non-response to anti-TNF therapy in CD, effective therapeutic alternatives have not yet been envisioned^14–18,24^. Here, we identify an IFNγ-dependent pathogenic mechanism associated with anti-TNF resistance that contributes to severe epithelial damage, with major implications for the clinical management. Patients with high IFIM/IRM ratios are less likely to benefit from anti-TNF Abs alone and are at increased risk of requiring surgical resection and postoperative disease recurrence. Importantly, the IFIM-associated program is targetable by upadacitinib, a JAK inhibitor approved for CD patients following anti-TNF therapy failure^75^. These findings provide a molecular rationale for identifying CD patients with active IFNγ-driven pathogenicity who may benefit from earlier introduction of JAK inhibitors within their treatment sequence.

The role of dysregulated mo-macs in supporting self-perpetuating chronic inflammation and disease progression in the gut is well-established but new strategies are needed to enhance our understanding of the molecular checkpoints associated with maladaptation of monocyte-to-macrophage trajectories in the CD inflamed ileum^14,15,33,36,37,44^. By decoupling the molecular programs associated with monocyte-to-macrophage maturation from those involved in functional and metabolic activities, our combinatorial analysis provides a granular resolution of the mo-mac compartment and its regulation in severe CD. In support of this, we observed that although inflammatory programs were detectable in cells with either monocyte or tissue macrophage features, the latter predominantly accounted for inflammatory activities in the healthy ileum. This aligns with the contribution of mucosal macrophages to the gut “physiological inflammation”, a concept originally proposed to explain the need for tonic host-microbiota interactions in order to maintain homeostasis in the steady sate^77^. Expanding on previous studies, including ours^15,36,50^, the present data strongly support a model in which most, if not all the disease-associated inflammatory tone in CD mo-macs arises from the accumulation of monocytes in inflamed mucosal areas. However, consistent with the high monocyte plasticity, we uncover a complex picture where this accumulation is marked by significant molecular heterogeneity. In the context of tissue injury, monocytes are rapidly recruited to the gut to serve inflammatory functions but also to promote return to homeostasis. Several models have been proposed to explain how monocyte contribute to the resolution of inflammation^78^. The “active repair” model suggests that mo-macs activate specific programs related to immunoregulation and tissue repair. Our results indicate that IRM simultaneously induced both proinflammatory and immunoregulation/repair programs, likely through the integration of NF-κB and TGFβ signaling pathways. This suggests that single monocytes may jointly engage in proinflammatory and pro-resolving activities, supporting the possibility of a cell-intrinsic mechanism for progressive functional reprogramming in response to the dynamics of microenvironmental cues. The “passive repair” model involves the gradual differentiation of monocytes into tissue macrophages^78^. While IRM retained expression of gene programs consistent with differentiation potential, we observed the highest levels of “transition” programs in Late_Mono, a group of monocytes lacking inflammatory activity. This implies that the mo-mac compartment in inflamed ileum of CD patients is partly comprised by a mixture of monocyte states engaged in both active and passive repair processes, potentially reflecting a physiological monocyte response aimed at resolving ongoing inflammation rather than driving disease severity.

In striking contrast, the program-based combinatorial strategy enabled us to characterize a distinct inflammatory monocyte state, which we refer to as IFIM. IFIM displayed unique inflammatory features and the absence of “Transition” and “Immunoregulation/Repair” programs. Enrichment of IFIM in CD inflamed ileums was associated with resistance to anti-TNF therapy and with severe clinical recurrence following initial ileocecal resection. IFIM, like IRM, activated inflammatory programs in response to NF-κB signaling but we uncovered a major role for IFNγ in shaping their molecular profile. This was evident not only in the strong interferon signature but, more importantly, in the unique induction of the “IFIM-program” program, which resulted from converging NF-κB and IFNγ pathway activities in monocytes. While previous works had suggested a role for IFNγ in inflammatory mo-macs in murine colitis and human IBD^79,23,80,81^, we now provide a comprehensive molecular cartography of the IFNγ-dependent and -independent inflammatory responses of monocytic cells in the inflamed ileum of patients with severe CD. Since its original identification as “macrophage activating factor”, IFNγ has been shown to extensively remodel the epigenome to alter gene transcription and drive mo-macs into a hyperinflammatory responsive state characterized by massive induction of inflammatory cytokines and NF-κB target genes^74^. Similarly, IFNγ induces a refractory state to anti-inflammatory factors and reverses so-called macrophage tolerance that follows prior NF-κB activation by microbial products and TNF. Importantly, IFNγ also interferes with the activity of transcriptional enhancers, part of which lose their ability to bind lineage-determining transcription factors PU.1 and C/EBP family proteins to stably target genes^82^. Our findings thus suggest that, beyond hyperinflammatory induction, monocyte exposure to IFNγ in the inflamed ileum directly contributes to prevent their maturation to a macrophage state. The localization of IFIM in inflamed ileums was strikingly confined to areas of severe epithelial damage. In non-responder patients to anti-TNF therapy, deep ulcerations have previously been associated with neutrophil accumulation driven by chemoattractants produced by inflammatory fibroblasts^17^. Our study reveals a complex cellular ecosystem in which IFIM cells, together with IFNγ-producing lymphocytes and iaDC, form a program recently recognized across diverse chronic inflammatory conditions^56^. Commensal and food-derived yeasts have recently been identified as antigenic drivers of IFNγ production by T cells with cytotoxic activity toward epithelial cells in the ileum of a subset of CD patients^83^. Together, this supports a scenario in which IFNγ-dependent IFIM and iaDC orchestrate a self-perpetuating cellular circuit at sites of ulceration, contributing to tissue damage and disease chronicity. This includes the chemokines CXCL9/CXCL10/CXCL11, which locally retain CXCR3⁺ lymphocytes, as well as IL-23, a cytokine secreted by monocytes in response to IFNγ and known to further amplify IFNγ production by T cells.

In conclusion, our findings provide a strong rationale for integrating molecularly informed patient stratification and monitoring to develop more effective, mechanism-guided therapeutic strategies. They also support approaches aimed at precisely reprogramming pathogenic monocyte trajectories in CD, rather than broadly targeting the mo-mac compartment, a strategy that has either proven ineffective or carried significant side effects due to the essential role of mo-macs in tissue homeostasis. These concepts have the potential to fundamentally redefine CD patient classification and may benefit the broader spectrum of IMIDs.

### Study limitations

While our *ex vivo* data show that upadacitinib rapidly suppresses IFIM-driven programs, establishing a direct link to clinical efficacy will require early, dynamic monitoring of these programs shortly after treatment initiation. It also remains to be tested prospectively whether IFIM activity can identify patients likely to resist anti-TNF therapy alone and guide rational sequencing of JAK inhibition to disrupt ulceration-site aggregates while preserving the benefits of anti-TNF therapy. In addition, prior clinical trials targeting IFNγ in CD demonstrated safety and efficacy only in a subset of patients, likely reflecting heterogeneous inclusion without molecular stratification for IFIM-driven pathology.

## MATERIALS AND METHODS

### Human subjects

Crohn’s disease (CD) patients eligible for inclusion in the study were identified by screening surgical programs at the CHU Nantes. Ileum CD tissues were obtained after first surgical resection. Protocols were reviewed and approved by the Institutional Review Board (IRB) at CHU Nantes (DC-2017-2987). Clinical characteristics, including Montreal classification, of the patients are summarized in Table S1. All macroscopically inflamed tissues included in the study were confirmed by pathological examination as active ileitis with transmural chronic inflammation.

### Blood collection

Blood Buffy Coat from healthy human volunteers was collected at Etablissement Français du Sang (EFS). In accordance with the Public Health Code (articles L 1121-1-1 et L 1121-1-2), a written consent and agreement from a person protection committee are not mandatory for non-interventional research in humans.

### Intestinal *lamina propria* cell isolation

Tissues were collected in ice-cold complete medium (RPMI 1640, 10% heat-inactivated FCS, 10mM HEPES, 1mM Sodium Pyruvate, 1X non-essential amino-acids, 2mM L-Glutamine, 100U/ml Penicillin, 0.1 mg/ml Streptomycin; all from Gibco), and processed within one hour after termination of the surgery. To limit biased enrichment of specific immune and stromal populations related to local variations in the intestinal micro-organization, we pooled 20 to 40 mucosal biopsies sampled all along the resected specimens using biopsy forceps (Fartley) to prepare cell suspensions. Epithelial cells were dissociated by incubating the biopsies in an EDTA-enriched dissociation medium (HBSS no calcium no magnesium (Gibco), 10mM HEPES (Gibco), 5mM EDTA (Sigma-Aldrich)) at 37°C under 100 rpm agitation for two cycles of 15 minutes. After each cycle, biopsies were hand-shaken for 30s and then vortexed vigorously for another 30s. Biopsies were then washed in complete medium, and transferred in digestion medium (HBSS calcium magnesium (Gibco), 2% FCS (Gibco), 0.5 mg/ml DNAse I (Sigma-Aldrich), 0.5 mg/ml Collagenase IV (Sigma-Aldrich)), for 40 minutes at 37°C under 100 rpm agitation. After digestion, the cell suspension was filtered through a 70µm cell strainer, and washed in PBS, 2% FCS, 2mM EDTA. Red blood cells were depleted using red blood cell lysis solution (distilled H_2_O, 155mM NH4Cl, 10mM KHCO3, 0.1mM EDTA, pH 7.2-7.4), and dead cells were depleted using the Dead cell removal kit (StemCell), following manufacturer’s recommendations. Viability of the final cell suspension was calculated using a hematocytometer and eosin exclusion and was routinely > 85%.

### Cell sorting

To enrich in intestinal MNP, cells obtained from the *lamina propria* were stained in PBS, 2% FCS, 2mM EDTA with a flow antibody cocktail (CD3, CD19, CD45, CD66b, CD326 and HLA-DR antibodies; all from BD Biosciences) for 20 minutes at 4°C in the dark. Cells were then washed, filtered through a 100µm cell strainer and stained with DAPI (Thermo Fisher scientific). Total cells (live CD326^-^ CD45^+/−^ cells) and MNPs (live CD326^-^ CD45^+^ CD66b^-^ CD3^-^ CD19^-^ HLA-DR^+^ cells) were FACS-sorted on a FACS Aria I cell sorter (BD Biosciences). Purity was routinely > 95%.

### Spectral flow cytometry

Cells obtained from the *lamina propria* were incubated with PBS containing purified NA/LE human Fc Block (BD Biosciences), Brilliant Stain Buffer Plus (BD Biosciences), ViaDye Red (Cytek Biosciences) and a panel of 23 antibodies for 30 minutes at 4°C in the dark (CD1c, CD3, CD11b, CD11c, CD14, CD16, CD19, CD24, CD38, CD45, CD64, CD66b, CD89, CD123, CD127, CD141, CD163, CD206, CD209, FcER1A, FOLR2, HLA-DR and PD-L1 antibodies; see Method Table for details). In vitro Momacs were stained with a reduced panel (PBS, Fc Block, Brilliant Stain Buffer Plus, ViaDye Red, and 5 antibodies: CD14, CD64, CD123, CD150, PD-L1). Cells were washed twice with PBS, 0.5% BSA, 2mM EDTA and acquired using a 5-laser Aurora spectral cytometer (Cytek Biosciences). Full emission spectra were determined with single-stained samples using compensation beads (ONEComp eBeads, Thermo Fischer Scientific). Cell autofluorescence was determined with unstained cells. Spectral deconvolution (or unmixing) algorithms were used from the SpectroFlo Software (Cytek Biosciences). Data were analyzed using Omiq software. Dead cells and doublets were excluded for all analyses.

### *Lamina propria* cell culture

Cells obtained from the *lamina propria* were resuspended at a concentration of 2 million cells per milliliter in complete medium (RPMI 1640, 10% heat-inactivated FCS, 10mM HEPES, 1mM Sodium Pyruvate, 1X Non-essential amino-acids, 2mM L-Glutamine, 100U/ml Penicillin, 0.1 mg/ml Streptomycin; all from Gibco). Cells were then cultured in a U-bottomed 96-well plate, with each well containing 400,000 cells. The cells were incubated for 40 hours with increasing doses of JAK-STAT inhibitor Upadacitinib (0.001, 0.01, 0.1, 1 and 10µM; from Eoromedex) or with DMSO as control. Supernatants were harvested at the end of the culture for cytokine quantification.

### Blood monocyte isolation and culture

CD14^+^ monocytes were isolated from Buffy Coats using StraightFrom Buffy Coat CD14 Microbeads (Miltenyi Biotec), following manufacturer’s recommendations. The positive fraction was eluted on a MultiMACS Cell24 Separator (Miltenyi Biotec). Red blood cells were depleted using red blood cell lysis solution (distilled H_2_O, 155mM NH4Cl, 10mM KHCO3, 0.1mM EDTA, pH 7.2-7.4) and platelets were depleted by spinning at 300g for 10 minutes. Purity of CD14^+^ monocytes was analyzed by flow cytometry and was routinely >98%. Monocytes were resuspended at a concentration of 1 million cells per milliliter in complete medium (RPMI 1640, 10% heat-inactivated FCS, 10mM HEPES, 1mM Sodium Pyruvate, 1X Non-essential amino-acids, 2mM L-Glutamine, 100U/ml Penicillin, 0.1 mg/ml Streptomycin; all from Gibco). Cells were then cultured in 24-well plates, with each well containing 1.5 million cells. The cells were incubated with different combinations of GM-CSF (1000U/mL), TNF (20ng/mL), IFNγ (10ng/mL); all from Miltenyi Biotec, and LPS (10ng/mL; Sigma-Aldrich). GM-CSF, TNF and LPS were added at day 0. Cells were washed at day 1 to remove LPS, and incubated with GM-CSF, TNF and IFNγ for the rest of the culture. In some experiments, JAK-STAT inhibitor Upadacitinib (Euromedex) was added at day 1, 30 minutes before adding IFNγ, and kept during the rest of the culture. Four concentrations of Upadacitinib were used (0.001, 0.01, 0.1 and 1µM), and mock conditions were treated with DMSO. At day 5, cell morphology was assessed by optical microscopy, and supernatants were collected for cytokine quantification. Cells were harvested by vigorous pipetting after incubation in PBS, 2% FCS, 2mM EDTA for 20 minutes at 4°C to help detaching the cells. When needed, the harvesting step was repeated until all the cells were recovered.

### Multiplex Immunoassays

Cytokine and chemokine levels in supernatants were quantified using a custom Luminex-based assay (Bio-Techne) enabling the detection of 24 analytes: CCL2, CCL3, CCL4, CCL7, CCL17, CCL18, CCL24, CXCL5, CXCL9, CXCL10, CXCL11, IL-1a, IL-1b, IL-1RA, IL-6, IL-8, IL-10, IL-12p70, IL-23, Total inhibin, M-CSF, Oncostatin M, Osteopontin, and S100A9. Cytokines were analyzed on a MagPix instrument and concentrations were expressed in pg/ml.

### RNA isolation

Total RNA was isolated using TRIzol reagent (Invitrogen), followed by RNeasy Mini Kit (Qiagen). RNA concentration and quality were determined using a QuBit Flex Fluorometer (Thermo Fiscsher Scientific) and Caliper LabChip GX (Life Sciences).

### Droplet-based single cell RNA sequencing

Cells were suspended at a concentration of 1 million cells per milliliter in PBS and 10,000 cells were processed with the Next GEM reagent kit v3 before being loaded onto the Chromium Controller instrument from 10x Genomics. Libraries were prepared using Chromium Single Cell 3′ Library Kits and sequenced on a NovaSeq 6000 (Illumina) with S1 flow cells. Base call files were converted to FASTQ and demultiplexed based on cell barcodes and unique molecular identifiers (UMI) and aligned to the GRCh38 human reference genome (Ensembl 98) using 10X Genomics Cell Ranger v5.0.0 to obtain a raw unfiltered matrix of counts.

### Public datasets

ScRNA-seq data of inflamed and uninflamed ileal samples from 11 patients with Crohn’s disease were downloaded from the NCBI Gene Expression Omnibus (GEO) public database under accession number GSE134809. ScRNA-seq data of 3 non-IBD controls, 2 uninflamed ileal and 12 inflamed ileal samples from treatment naïve adult Crohn patients in the TAURUS cohort were extracted from the processed data available on Zenodo (https://doi.org/10.5281/zenodo.13768607). Bulk RNA-seq data of the RISK cohort, composed of samples from 199 ileal biopsies from treatment-naïve pediatric Crohn patients and 15 non-IBD controls, were downloaded from GEO under the accession number GSE1348811. Microarray data of the REMIND cohort, composed of samples from 196 ileal biopsies of adult at the time of their first surgery and 25 non-inflammatory control, were downloaded from GEO under the accession number GSE186582.

### Generation of metacells and preprocessing

Raw unfiltered matrices from either the MNP-enriched samples (n=7) or the total lamina propria samples (n=52) were merged in a single matrix each, keeping only the common genes expressed across samples. Cells were then processed with the Metacells-2 algorithm^3^ (v0.8.0) on Python v3.10.4. Cells comprising less than 800 UMI or above 25000 UMI were excluded from the analysis based on inspection of per-sample UMI and gene count distribution. To improve the metacell generation, cells with contaminant genes were excluded: cells above 20% of hemoglobin genes (*HBA1, HBA2, HBB*) as a marker of red blood cell contamination and mitochondrial genes (genes starting by “MT-”, “MT1” or “MTRNR”) as a marker of apoptotic or lysing cells. Given the immune-focused scope of this study, cells expressing more than 10% of epithelial genes were filtered-out as well. Epithelial gene list: *PLA2G2A, CLCA1, REG4, S100A14, ITLN1, ELF3, PIGR, EPCAM, REG1B, REG1A, REG3A, FABP1, RBP2, SST, FABP2, SPINK1, FABP6, AGR2, AGR3, CLDN3, CLDN4, DEFA6, DEFA5, SPINK4, ALDOB, LCN2, MUC2, KRT8, KRT18, TSPAN8, OLFM4, GPX2, IFI27, PHGR1, MT1G, CLDN7, KRT19, FXYD3, LGALS4, FCGBP, TFF3, TFF1.* Additional genes like the long noncoding RNA (lncRNA) MALAT1 and XIST, ribosomal genes, as well as Immunoglobulin chain genes, and HLA class I and II genes, were excluded to prevent highly variable but biologically uninformative genes from dominating metacell construction across patients. A target metacell size of 40000 UMIs was chosen for the global metacell object. Based on the gene marker analysis of known cell type markers, lineages were segregated to recompute more precise metacells to have a more comprehensive representation of the different cell states. The target metacell size was adapted depending on the number of cells and was reduce down to 5000 UMIs for the Macrophage and DC metacells objects. From 14386 cells in the macrophage dataset and 4144 cells in the DC dataset, we obtained respectively 802 and 202 metacells. The feature count in metacells is defined as a weighted arithmetic mean of the genes in the cells pooled in a metacell or in the case of noisy genes, a normalized geometric mean. The cell sizes (total UMI counts per cell) are also considered so that large cells don’t dominate the average, as per the metacells package documentation.

### *In silico* selection of mo-macs in MNP-enriched suspensions

Flow-sorted mononuclear phagocytes (MNPs) from lamina propria cells (**Figure S1A**) were freshly isolated from inflamed mucosae of patients with stricturing ileal disease at time of first surgical resection and analyzed by single-cell RNA-sequencing (scRNA-seq) using Metacells-2^52,53^. MNP metacells (*LYZ*, *HLA-DRA*) were separated into mo-macs (*S100A9, C5AR1*, *FCG3RA*, *MAF*, *C1Qs*, *CD209*) and dendritic cells (DCS; *FLT3*, *ZBTB46*, *LY75*), including subsets of DC1 (*XCR1*, *CLEC9A*, *CADM1*), DC2/3 (*CLEC10A*, *FCER1A*, *CD1C*) and activated DCs (*LAMP3*, *CD200*, *CD274*). Plasmacytoid dendritic cells (pDCs; *GZMB*, *TCF4*, *IL3RA*, *LILRB4*), along with minor contaminating populations of plasma cells (*TNFRSF17*), B cells (*MS4A1*) and fibroblasts (*LUM*, *PDGFRB*) were also identified (**Figure S1B**). Mo-mac metacells were selected for further analysis using a gene module-based approach.

### Gene module identification

The rationale behind the selection of gene modules is that highly correlated gene expression patterns might indicate a functional relationship and participation in a common molecular pathway. By organizing genes into modules, we aimed to capture specific and shared functional pathways within the dataset. The first step of the module identification is the selection of genes based on their expression levels and variability in metacells. These variable genes are selected using a loess curve fit on the log(variance/mean) versus log(mean) distribution as performed in Martin et al^1^ and adapted by lineages. We first computed the pairwise Pearson correlations between all the variable genes to identify groups of co-expressed genes. We then performed hierarchical clustering to organize the co-expression matrix into 150 distinct groups called modules. A signature score *S_s_* is defined as the sum of the log-transformed gene signature expression in a metacell normalized by the total expression of this given metacell.

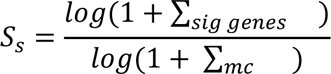

A z-transformation was then applied to compare signature enrichment across metacells or clusters of metacells. Mo-mac metacells were grouped by hierarchical clustering, which allowed to curate 17 molecular programs, biologically annotated based on gene composition and/or the Molecular Signature Database (MSigDB)^54^.

### Gene set enrichment analysis

Gene programs were compared to MSigDB Hallmark gene sets to identify significant overlaps (www.gsea-msigdb.org/gsea/msigdb/annotate.jsp). The queried gene sets were obtained from version v2023.1.Hs of MSigDB^4^.

### Annotation of mo-mac programs

Program VII, VIII and V were respectively annotated as “MonoA”, “MonoB” and “Macrophage” based on the enrichment of genes associated with monocyte identity (MonoA: *SELL*, *CD55*, *FCAR*, *RETN*; MonoB: *S100A8/A9*, *S100A12*, *VCAN*, *FCN1*) and macrophage identity (Macrophage: *C1Qs*, *CD209*, *MERTK*, *FUCA1, SELENOP*) in the gut^35,45,48^. Program XIII and XIV were annotated as “Inflammatory_A” (*IL23A*, *IL6*, *CXCL2*, *CXCL3*, *CCL20*, *NLRP3*, *ICAM1*, *CD40*, *CD83*, *SLAMF7*, *PLAUR*) and “Inflammatory_B” (*IL1B*, *TNF*, *CCL3*, *CCL4*, *CXCL8*, *F3*) based on gene composition and significant overlaps significantly MSigDB Hallmark “TNFA Signaling via NFKB” and “Inflammatory Response” (adjusted P <10^^-10^ for both). Program XV was annotated as “Inflammatory_Mono1” because of its specific detection in Mono1 and significant overlap with “Hallmark Inflammatory Response” (adjusted P <10^^-10^). Program XII was annotated as “Immunoregulation/Repair” based on its composition enriched in genes involved in immunoregulation (*IL10*, *NFKBID, OLR1*)^58^, epithelial wound healing and tissue repair (*EREG*, *MMP19*)^59^. Program IX was annotated as “Interferon” (*IRF1*, *CXCL10*, *IFIH1*, *GPB4*, *WARS1*, *CIITA*, *OAS2*) based on gene content and overlap with “Hallmark Interferon Response” (adjusted P <10^^-10^). Programs I, IV and VI were annotated as “Transition” because of gene content and their upregulation in Mono5, a group we considered as late maturation stage monocytes as compared to early Mono4. Program II was annotated as “OXPHOS” because of significant overlaps with “Hallmark Oxidative_Phosphorylation, “Hallmark Myc_Targets” and “Hallmark Fatty_Acid_Metabolism” (adjusted P <10^^-10^ for all three). Program XVI was annotated as Hypoxia due to overlaps with “Hallmark Hypoxia” (adjusted P <10^^-10^) and gene content linked to glycolysis (*SLC2A1*, *LDHA*, *HK2*, *ERO1A*). Program XVII was annotated as stress because of significant overlap with “Hallmark HSF” (adjusted P <10^^-10^).

### RNA-seq analysis

The count matrix for RISK dataset represents fragments per kilobase of transcript per million (FPKM) estimates. FPKM estimates where transformed using log(1+FPKM) before calculating enrichment scores per sample, defined as the sum of the log-transformed gene signature expression in a sample normalized by the total expression of this given sample. The microarray data obtained for the REMIND cohort being already normalized, the program enrichment scores were directly computed on the normalized expression matrix. Bulk RNA-seq data obtained from blood samples were filtered to exclude failed experiments and samples with low read counts. Data were normalized in counts per million (CPM) using the cpm function from the edgeR R package with default parameters and transformed using log(1+CPM). Program enrichment scores were then computed as previously described.

### PROGENY

PROGENy (Pathway RespOnsive GENes) provides gene programs reflecting the downstream effects of pathway activation or inhibition, derived from a large collection of manually-curated perturbation experiments (Schubert et al., 2018). Using the progeny R package (version 1.17.3), 500 top genes associated with selected pathways are extracted from the full human model. Each gene has an associated weight to the pathway, for which a positive weight translates to the gene up-regulation during the pathway signaling. Genes with a negative weight are therefore filtered out to focus on genes induced or up-regulated by the given pathway, before computing the program enrichment score as previously described in “Gene module identification”.

### Motif enrichment analysis

The motif enrichment analysis was performed with the RcisTarget R package (version 1.10.0, Verfaillie et al., 2015), using the human motif database hg38 (version 9, https://resources.aertslab.org/cistarget/databases/homo_sapiens/hg38/refseq_r80/mc9nr/gene_based/). The database ranking the motif-gene association scores used in our analyses includes both the close promoter region of 500 base-pair (bp) upstream to 100 bp downstream of the transcription starting site (TSS) and the extended search space of 10 kilobases up and down the TSS. To explore potential transcription factor enrichment, we used SCENIC (version 1.3.1, Aibar et al., 2017) which aggregates individual regulons based on gene expression with GRN Boost and filters genes through the RcisTarget calculation of normalized enrichment score (NES). NES value determines genes enriched in the TF binding motif in their promoter region (NES > 3). The AUCell step was ignored due to unsatisfactory automatic selection.

### Ligand Receptor analysis

Interactions tested in this analysis are based on the CellPhoneDB database resource of hand-curated ligand-receptor pairs (CellPhoneDB v5, Troulé et al., 2025) and an additional selection from experimentally validated pairs (Ramilowski et al., 2015). This list was first restricted to the protein-protein interactions associated to the “Ligand-Receptor” directionality and for whom both ligands and receptors are expressed in our data. Two categories of patients were defined in the CD inflamed ileum group, based on the proportion of Mono1 in the monocyte compartment. Samples with ≥ 50% of Mono1 are called Mono1^hi^ (patients 128, 138, 181, 187, GIM31, GIM33, GIM35, GIM36 and GIM38) while samples below this threshold are called Mono1^low^ (patients 69, 158, 190, 193, 196, 209, GIM7, GIM8, GIM21 and GIM23). Pairs intensity scores and statistical power were defined as previously described by Martin et al. (2019). A Ligand-receptor pair intensity is the product of the globally normalized ligand gene expression in monocytes and the normalized receptor expression in the selected lineage, for each of the two patient groups. The color values represent the log10 difference in interaction intensity scores between the two patient groups. Cells from the two groups are shuffled for 105 permutations and the p-values are then calculated as the proportion of permutations where the absolute log2 fold change exceeded the observed value, followed by a Benjamini-Hochberg correction for multiple testing.

### ScRNA-seq-based inferred secretome in monocytes

Genes in the inferred secretome of monocytes were selected from all enriched ligands of the ligand:receptor analysis, and filtered manually or based on their differential expression across monocyte subtypes.

### Xenium sample preparation

Samples and data were processed as described in Thomas J et al. (https://doi.org/10.1101/2025.05.19.654443). FFPE-preserved tissues were sectioned onto Xenium slides and processed for deparaffinization and decrosslinking following the 10x Genomics FFPE protocol (CG000580). The Xenium slides were then processed according to the 10X Genomics Xenium In Situ Gene Expression user guide (CG000582) for probe hybridization, ligation and amplification. Tissue slides were incubated with a customized gene panel comprising the 377 genes from the Xenium Multi-Tissues and Cancer panel and 100 additional genes (Table S9). The custom genes were curated from the literature and identified as cluster-enriched in scRNA-seq data to optimize population resolution, with a focus on MNP (46 identity/state-defining genes), as well as 35 functional genes and 19 identity/state-defining genes of other lineages. Panel composition was iteratively benchmarked against scRNA-seq datasets using prediction-based classification to ensure accurate resolution of MNP subpopulations as described in Thomas J et al. Following these steps, slides were loaded into the Xenium Analyzer and acquired.

### Xenium data processing

Xenium Analyzer output are raw multi-cycle fluorescence images that are first aligned and corrected to create spatially consistent image stacks. These image stacks were then analyzed to find the rolling circle amplification (RCA) products, as localized fluorescent spots that corresponds to the transcripts captured in the tissue. The signal intensities of these RCA products were interpreted using a Xenium codebook that assigns an expected pattern of fluorescence signal to a gene or a negative control probe. Each decoded transcript was assigned a quality score (Q-score) reflecting the confidence in the transcript identity. The transcripts were then assigned to their cells by using the probabilistic segmentation method Proseg (Jones et al., 2024) with default parameters. These count matrices were analyzed using the scanpy and squidpy python packages, and log-normalized using the pp.normalize_total and pp.log1p functions. A total of 1,802,727 cells were identified across the 8 samples. Cell annotations were performed in a gating-like fashion, by sequentially identifying celltypes and lineages with specific reduced gene signatures. We first identified Epithelial cells (*EPCAM, AGR3, GPX2, MMP7*), Endothelial cells (*VWF, SELE, ACKR1*), Lymphatic stromal cells (*LYVE1, MMRN1, PROX1*), Glial cells (*LGI4, SOX2, PCSK2*), Mast cells (*MS4A2, CPA3, CTSG*), and then more broad lineages: B cells (*MS4A1, BANK1, CD19*), T and ILC cells (*CD3D, CD2, CD3E, IL7R, KIT, IL23R*), MNPs and Neutrophils (*CD68, HLA-DRB1, CD14, CD4, CXCR2, MME, ITGAX, LILRB4*), Fibroblasts (*PDGFRA, PDGFRB, PTGDS, ACTA2, MYH11*), Plasma cells (*TNFRSF17, DERL3, TENT5C, MZB1*).

### Statistical analysis

All statistical analyses were performed using R (version 4.0.3) language for sequencing data and in Graphpad Prism (version 7.0) for cytometry data.

Comparisons of cell distributions between multiple patient groups were performed using a two-sided Kruskal–Wallis test, followed by Dunn’s post hoc pairwise comparisons with Holm’s correction for multiple testing. Comparisons of cell distributions or gene programs between two patient groups were performed using Mann-Whitney *U* tests. Gene expression between mo-mac subtypes were conducted using pairwise Mann-Whitney *U* tests, with Bonferroni correction applied to adjust for multiple testing when appropriate. Program enrichment was evaluated by comparing the median enrichment score of each mo-mac subtype against the global mean using Wilcoxon rank-sum tests, followed by Bonferroni correction for multiple comparisons. Pairwise associations between subtypes and IFIM/IRM distributions were assessed using Spearman’s rank-order correlation coefficient. For cytometry analyses, differences in PDL1+CD123+ frequencies and cytokine levels across increasing concentration of upadacitinib concentration were assessed using Friedman tests, with Dunn’s post-hoc multiple comparison tests between all conditions. For each test performed, p-values lower than 0.05 were considered statistically significant.

## DATA AND CODE AVAILABILITY

FASTQ sequencing files and processed count matrices for single-cell and bulk RNA sequencing will be made available on GEO upon publication. The R code and shiny app to reproduce analyses and figures shown in this study will be available on GitHub (https://github.com/Mr-Laurent/laurent_et_al_2025) upon publication.

Spatial transcriptomics data were obtained from Thomas J et al. and will be publicly available on GEO upon publication. (https://doi.org/10.1101/2025.05.19.654443).

## ACKNOWLEDGMENTS

We sincerely thank all the patients who participated in this study. JCM received support from NExT “Junior Talent” and ANR JCJC (ANR-20-CE17-0009). LOL is supported by a doctorate fellowship from fondation pour la recherche médicale (FRM). NC received a doctorate fellowship from fondation pour la recherche médicale (FRM). The authors thank the biological resource centre for biobanking (CHU Nantes, Hôtel Dieu, Tumorothèque, Nantes, F-44093, France.

## AUTHOR CONTRIBUTION

Conceptualization, JCM.; Data Curation, TL, MM, GB; Formal Analysis, TL, LOL, MB; Investigation, TL, MM, JRT, GB, LB, AJ, CB, JC, CF, NC, TS, CG, JFM, CLB, JP, EDD, AB, ASM, AS; Methodology, TL, JRT, GB, MU, PCH, EK, JP, AS; Supervision, J.C.M; Visualization, N.D., J.C.M.; Writing – Original Draft, TL, MM, JCM.; Writing – Review & Editing, JFC, SM, EK, JCM.

## CONFLICT OF INTEREST DISCLOSURES

**Figure S1:**
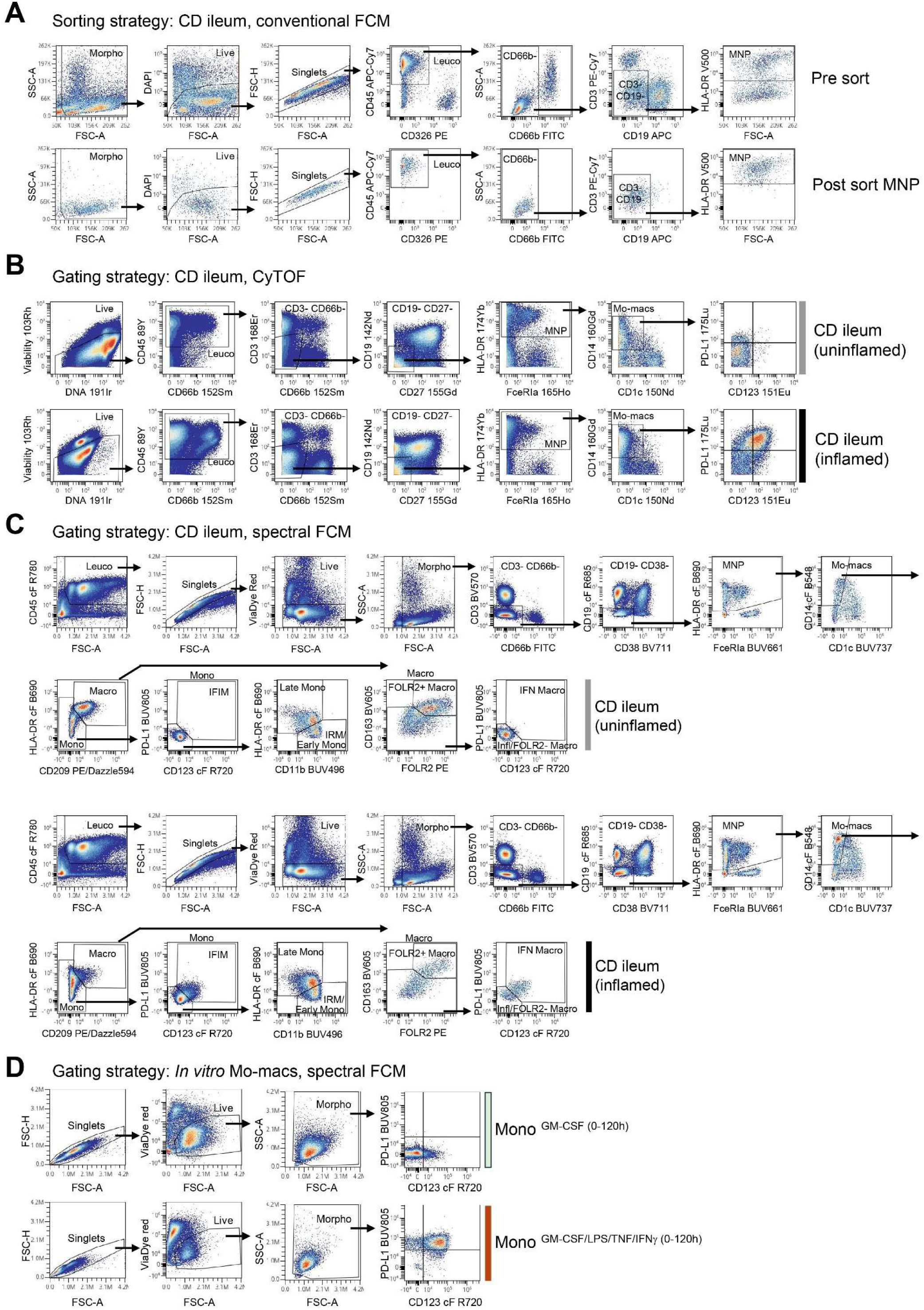
Representative gating strategies for human MNP. **(A)** Representative sorting strategy for *lamina propria* MNP analyzed by conventional flow cytometry. Total cells from inflamed CD ileums pre-sort (top) and sorted MNPs (bottom) are shown. **(B)** Representative dot plots of the step-by-step gating strategy to define PD-L1^+^ CD123^+^ cells in total mo-macs in the Mount Sinai cohort analyzed by CyTOF. Uninflamed (top row) and inflamed (bottom row) CD ileums are shown. **(C)** Representative dot plots of the step-by-step gating strategy to define FCM_mo-mac groups in the CHU Nantes cohort analyzed by spectral flow cytometry. Uninflamed (top row) and inflamed (bottom row) CD ileums are shown. **(D)** Representative gating strategy (spectral flow cytometry) for blood monocytes cultured 5 days with GM-CSF (top) or GM-CSF, LPS, TNF and IFNγ (bottom).

**Figure S2:**
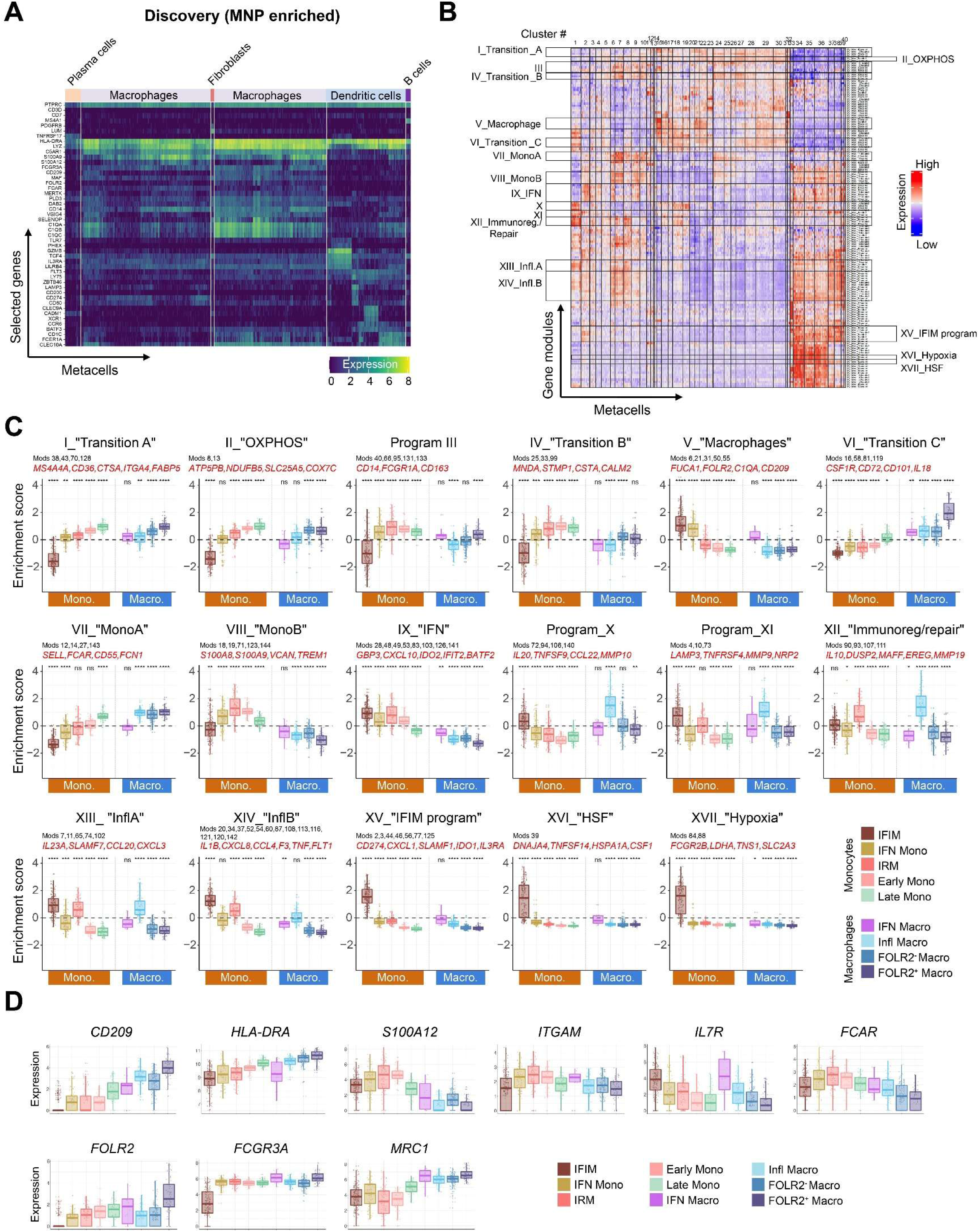
Mo-mac molecular programs in CD ileums. **(A)** Heatmap showing expression of representative genes (columns) in single metacells (rows) grouped as indicated. **(B)** Heatmap showing the expression of 150 gene modules across 802 mo-mac metacells grouped into 40 clusters. The analysis of module expression patterns and gene composition allowed to defined 17 molecular programs. **(C)** Boxplots comparing expression levels of indicated programs across mo-mac groups. Gene modules included in the program are indicated. In red are selected representative genes. P-values were calculated using Wilcoxon signed-rank tests (group median vs. global mean, **Table S21**) with Bonferroni correction between all groups. *\*P < 0.05, **P < 0.01, ***P < 0.001, ****P < 0.0001, n.s. non-significant.* **(D)** Boxplots comparing expression levels of indicated genes across mo-mac groups.

**Figure S3:**
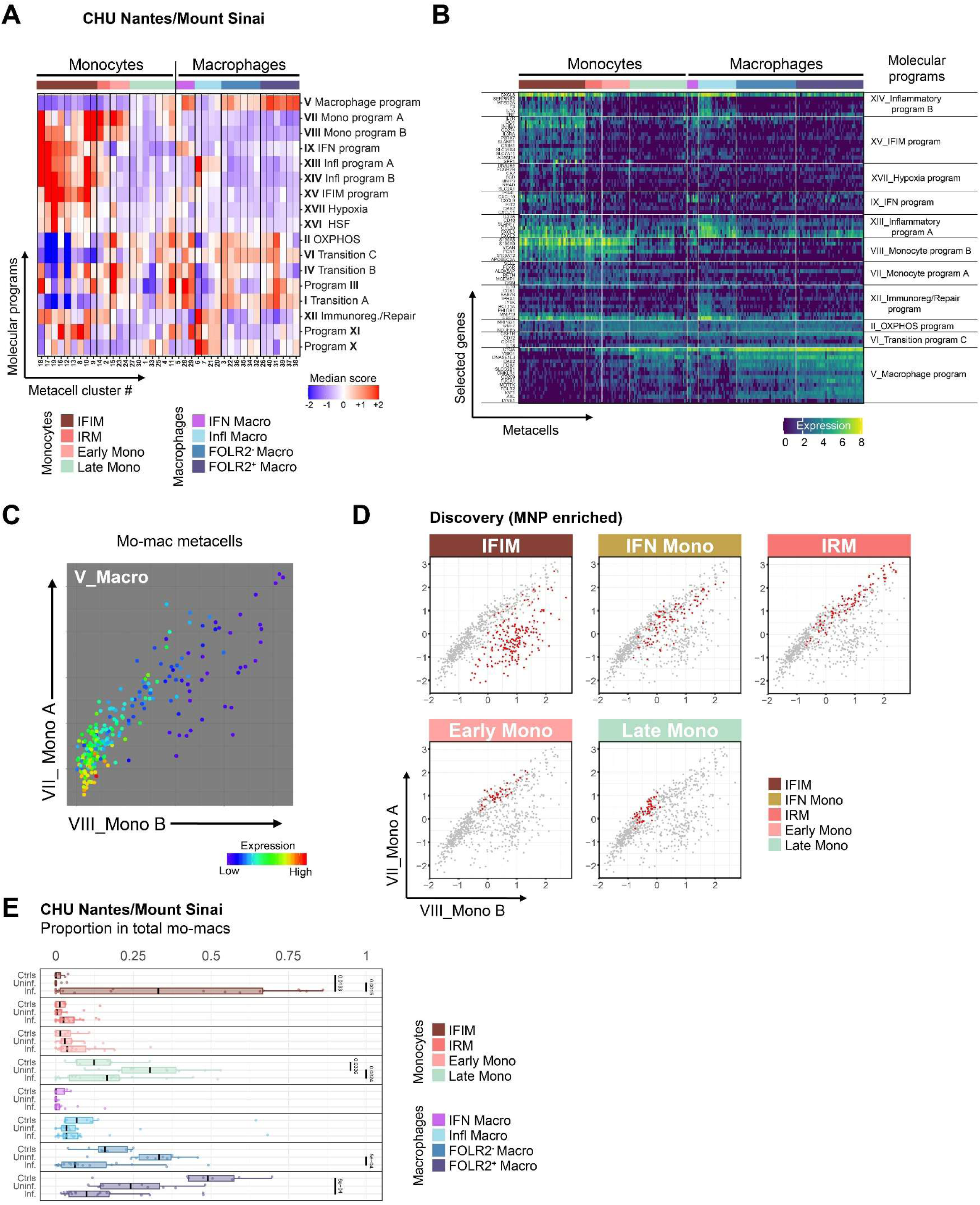
A state of inflammatory IFIM accumulates in CD inflamed ileum. Mo-macs were identified in total *lamina propria* cells from patients in the CHU Nantes/Mount Sinai cohort (control ileums n=8; control colons n=4; CD uninflamed ileums n=9; CD inflamed ileums n=18; UC inflamed colons n=4). **(A)** Heatmap showing expression of the 17 molecular mo-mac programs (rows) in metacell clusters (columns) grouped according to combinatorial profiles into 8 molecular states of mo-macs. IFN_Mono were not identified in this analysis. **(B)** Heatmap showing expression of representative genes in mo-mac programs (rows) in single metacell (columns) grouped in mo-mac states as defined in A. **(C)** Scatterplot of mo-macs metacells according to their expression of the two monocyte gene programs (Monocyte program A - y-axis; Monocyte program B – x-axis) and the macrophage program (Macrophage program – color-coded). **(D)** Scatterplots of mo-macs metacells according to their expression of the two monocyte gene programs. In red are the metacells annotated in the group of monocytes indicated on top of each scatterplot. **(E)** Boxplots showing the proportion of mo-mac groups in total mo-macs among indicated groups. Each dot represents a single ileum sample, and lines represent median and quartiles. The whiskers represent 1.5 times the interquartile range. P-values were obtained by Dunn’s post hoc test following Kruskal–Wallis, adjusted with Holm correction for multiple testing (**Table S22**).

**Figure S4:**
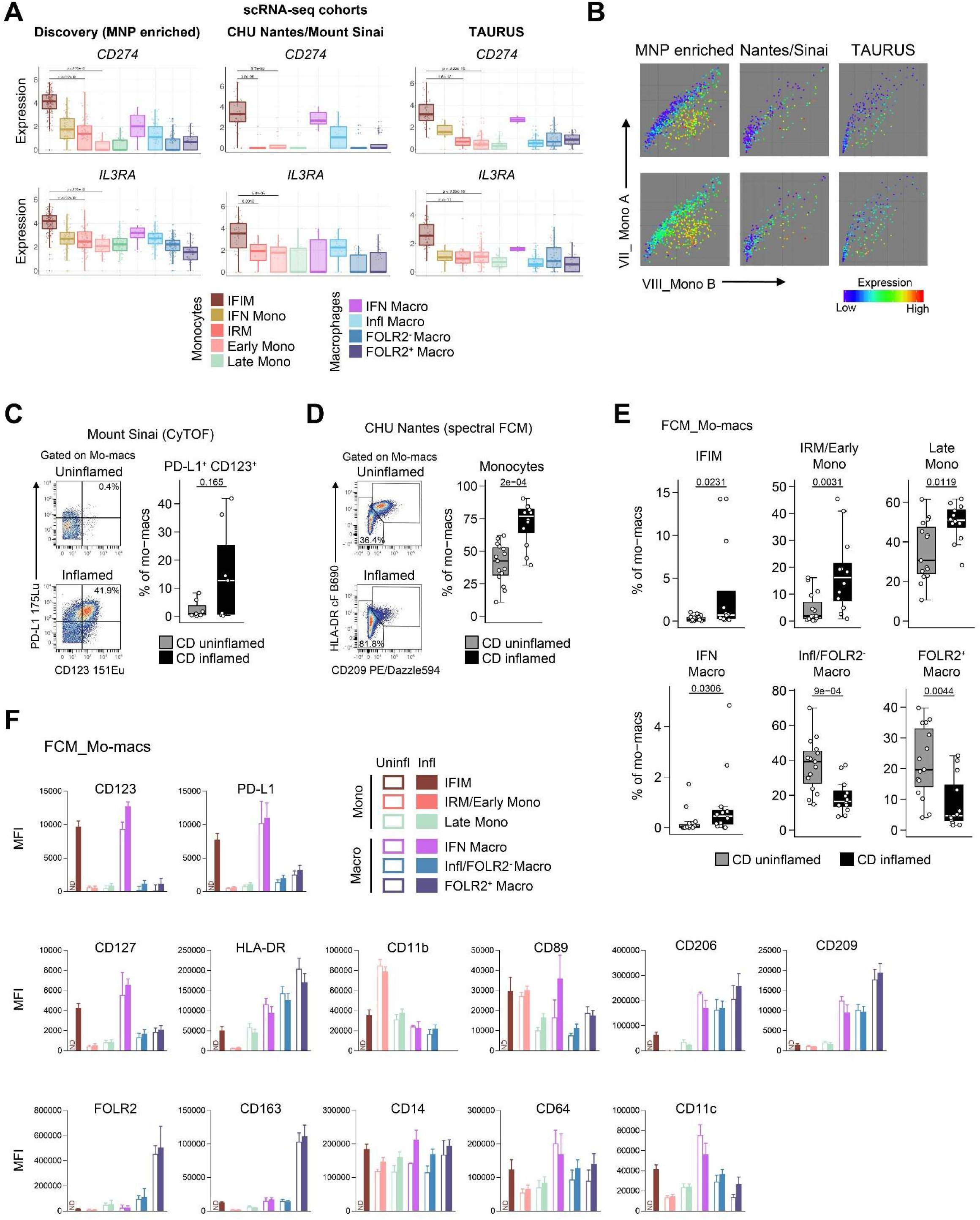
Phenotypic characterization of mo-mac states in CD ileum. **(A)** Boxplots comparing expression levels of indicated genes across mo-mac groups in mo-macs from MNP-enriched suspensions (left); the CHU Nantes/Mount Sinai cohort (middle) and the TAURUS cohort (right). P-values were calculated using a Mann-Whitney *U* test with Bonferroni correction. For clarity, only relevant comparisons are shown. **(B)** Scatterplots of mo-macs metacells according to their expression of the two monocyte gene programs (Monocyte program A - y-axis; Monocyte program B – x-axis) and the color-coded indicated single genes in MNP-enriched suspensions (left); the CHU Nantes/Mount Sinai cohort (middle) and the TAURUS cohort (right). **(C)** Representative FACS plots and bar plots (mean +/− s.e.m.) showing the proportions of PD-L1+ CD123+ cells in total mo-macs in the Mount Sinai cohort analyzed by CyTOF. **(D)** Representative FACS plots and bar plots (mean +/− s.e.m.) showing the proportion of monocytes in total mo-macs in the CHU Nantes cohort analyzed by spectral flow cytometry. P-values were calculated using a Mann-Whitney *U* test (**Table S23**). **(E)** Box plots comparing the proportions of indicated FCM_mo-mac groups between uninflamed and inflamed CD ileums. Each dot represents an ileum sample. P-values were calculated using a Mann-Whitney *U* test (**Table S23**). **(F)** Bar plots (mean +/− s.e.m.) of MFI levels of the indicated markers across mo-mac groups from uninflamed (light colors) and inflamed (dark colors) CD ileums. ND. not detected.

**Figure S5:**
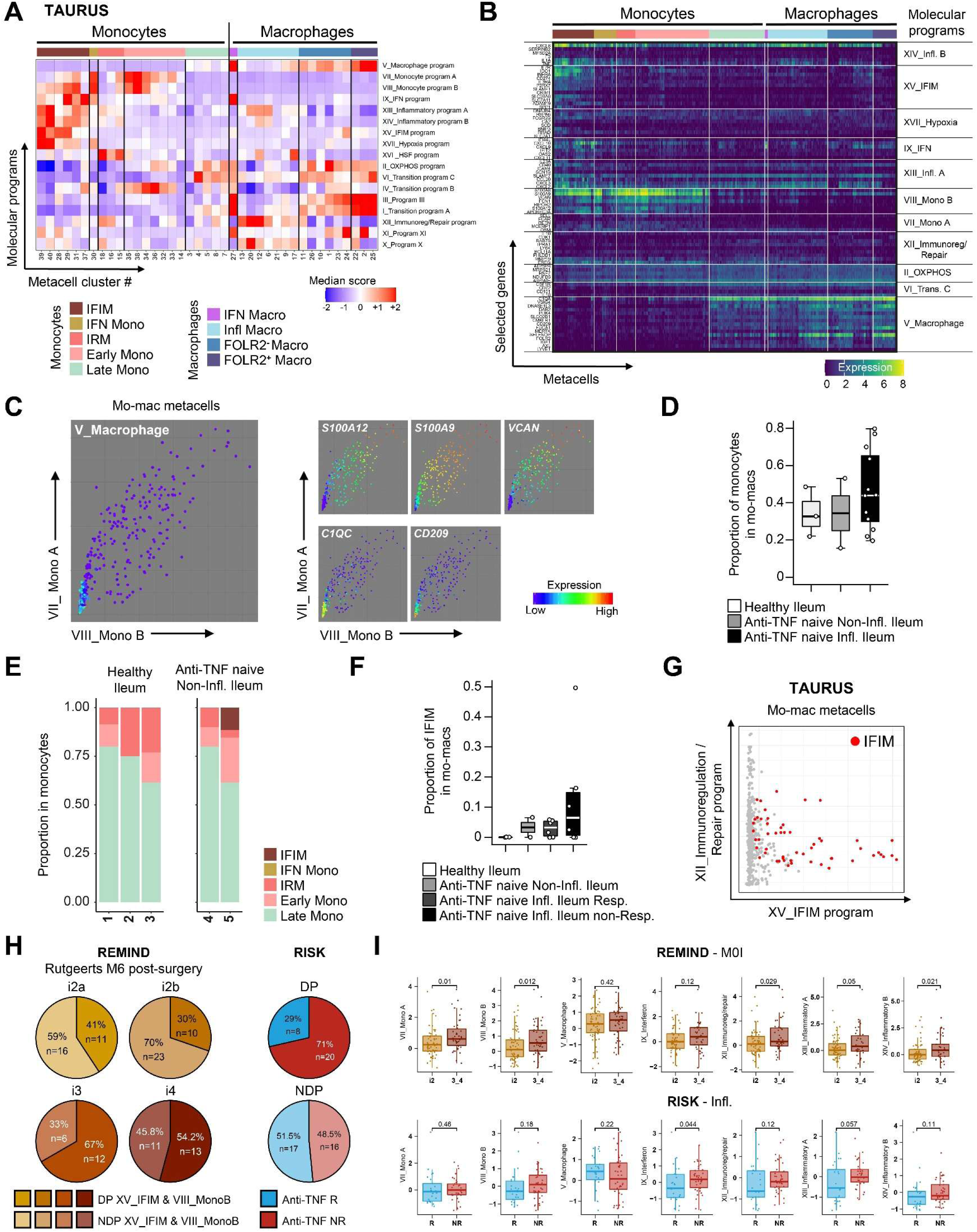
Clinical associations of program-based monocyte profiles in CD. Mo-macs were identified in total *lamina propria* cells from patients in the TAURUS cohort. **(A)** Heatmap showing expression of the 17 molecular mo-mac programs (columns) in metacell clusters (rows) grouped according to combinatorial profiles into nine molecular states of mo-macs. **(B)** Heatmap showing expression of representative genes in mo-mac programs (rows) in single metacell (columns) grouped in mo-mac states as defined in (A). **(C)** Scatterplots of mo-macs metacells according to their expression of the two monocyte gene programs (MonoA - y-axis; MonoB – x-axis) and the macrophage program (Macrophage – color-coded) (left) or indicated single genes (right). **(D)** Box plots comparing the proportions of monocytes in total mo-mac in the ileum between indicated groups. Each dot represents a CD patient, and lines represent median and quartiles. The whiskers represent 1.5 times the interquartile range (IQR, **Table S27**). **(E)** Bar graphs showing monocyte profiles across patients. Each bar corresponds to one patient in the TAURUS cohort in the healthy and uninflamed conditions **(F)** Boxplots comparing the proportions of IFIM in total mo-macs in the ileum between indicated groups. Each dot represents a CD patient, and lines represent median and quartiles. The whiskers represent 1.5 times IQR (**Table S27**). **(G)** Scatterplots of total mo-mac metacells according to their expression of programs “Immunoregulation/Repair” (y-axis) and “IFIM-program” (x-axis). **(H)** Pie charts showing the proportion of patients co-expressing (dark colors) or not co-expressing (light colors) mo-macs programs “MonoB” and “IFIM-program” in the REMIND cohort according to their Rutgeerts score 6-months post-surgery (left). Pie charts showing the proportion of future anti-TNF non-responders or responders among patients co-expressing (dark colors) or not co-expressing (light colors) mo-macs programs “MonoB” and “IFIM-program” in the RISK cohort (right). **(I)** Boxplots showing the expression of indicated programs between indicated groups in the REMIND (top) and RISK (bottom) cohorts. Each dot represents a single ileum sample, and lines represent median and quartiles. The whiskers represent 1.5 times the interquartile range. P-values were calculated using Mann-Whitney *U* tests (**Table S25**).

**Figure S6:**
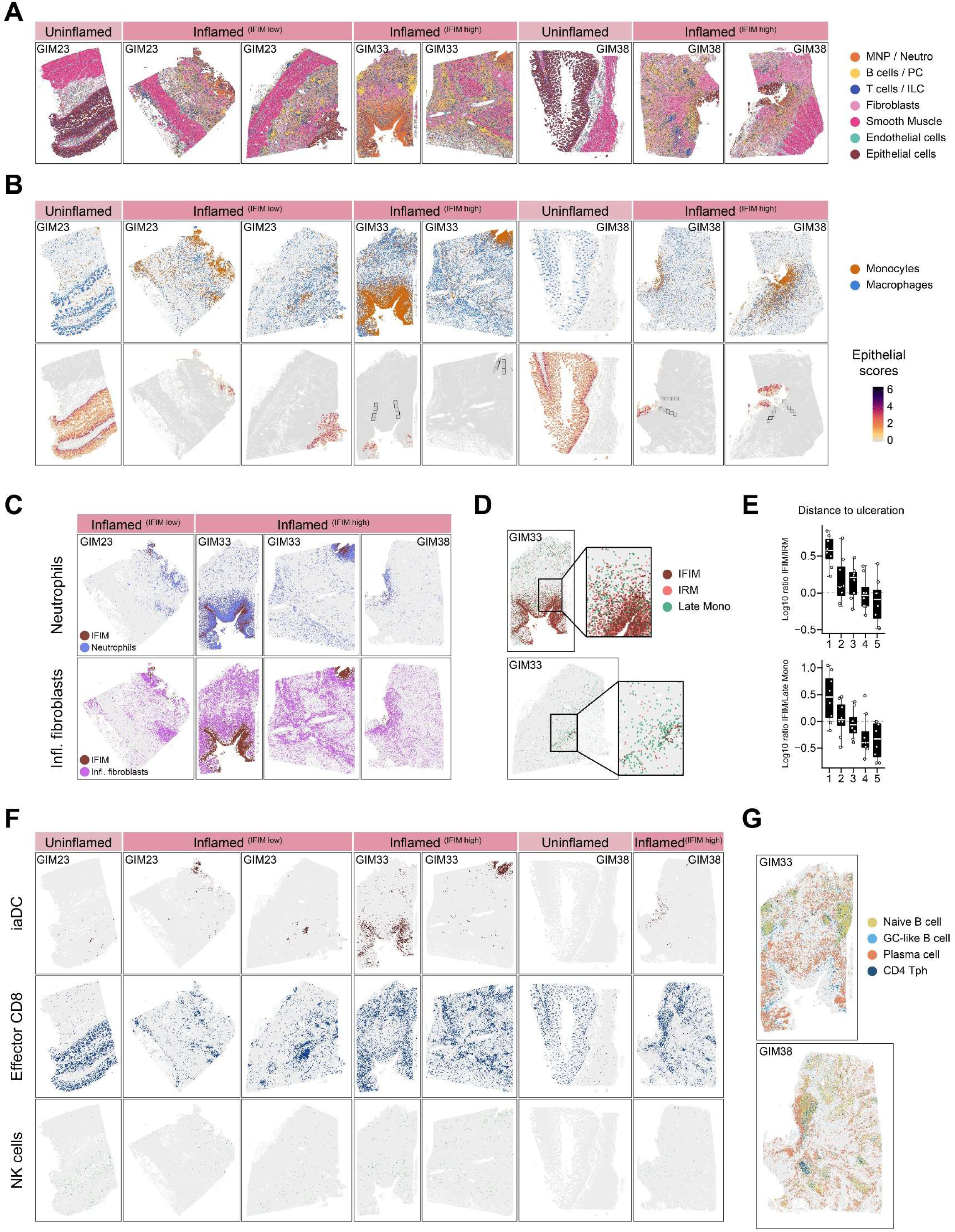
Enrichment in inflammatory IFIM at sites of severe epithelial damage. **(A)** Spatial visualization of main lineages localization in uninflamed and inflamed ileum cross-sections **(B)** Spatial visualization of monocytes and macrophages localization in uninflamed and inflamed ileum cross-sections (top). Black squares outline the selected zones of ulceration to deeper tissue gradients in inflamed ileums. Quantification of an epithelium gene score (bottom) highlights zones of epithelial damages in inflamed ileums. **(C)** Spatial visualization of neutrophils (top, blue) and inflammatory fibroblasts (bottom, purple) distribution in inflamed ileum cross-sections, with IFIM position showed in brown. **(D)** Spatial visualization of the monocyte subtype distribution in the inflamed ileum of IFIM enriched patient (GIM33) and IFIM low patient (GIM23) **(E)** Boxplots comparing the IFIM/IRM (top) and IFIM/Late Mono ratio (bottom) across the selected zones in inflamed ileum cross-sections. **(F)** Spatial visualization of iaDC (top row), effector CD8 Tcells (middle row) and NK cells (bottom row) in uninflamed and inflamed ileum cross-sections. **(G)** Spatial visualization of the main cell subtypes constituting the TLS structures in inflamed ileum cross-sections.

**Figure S7:**
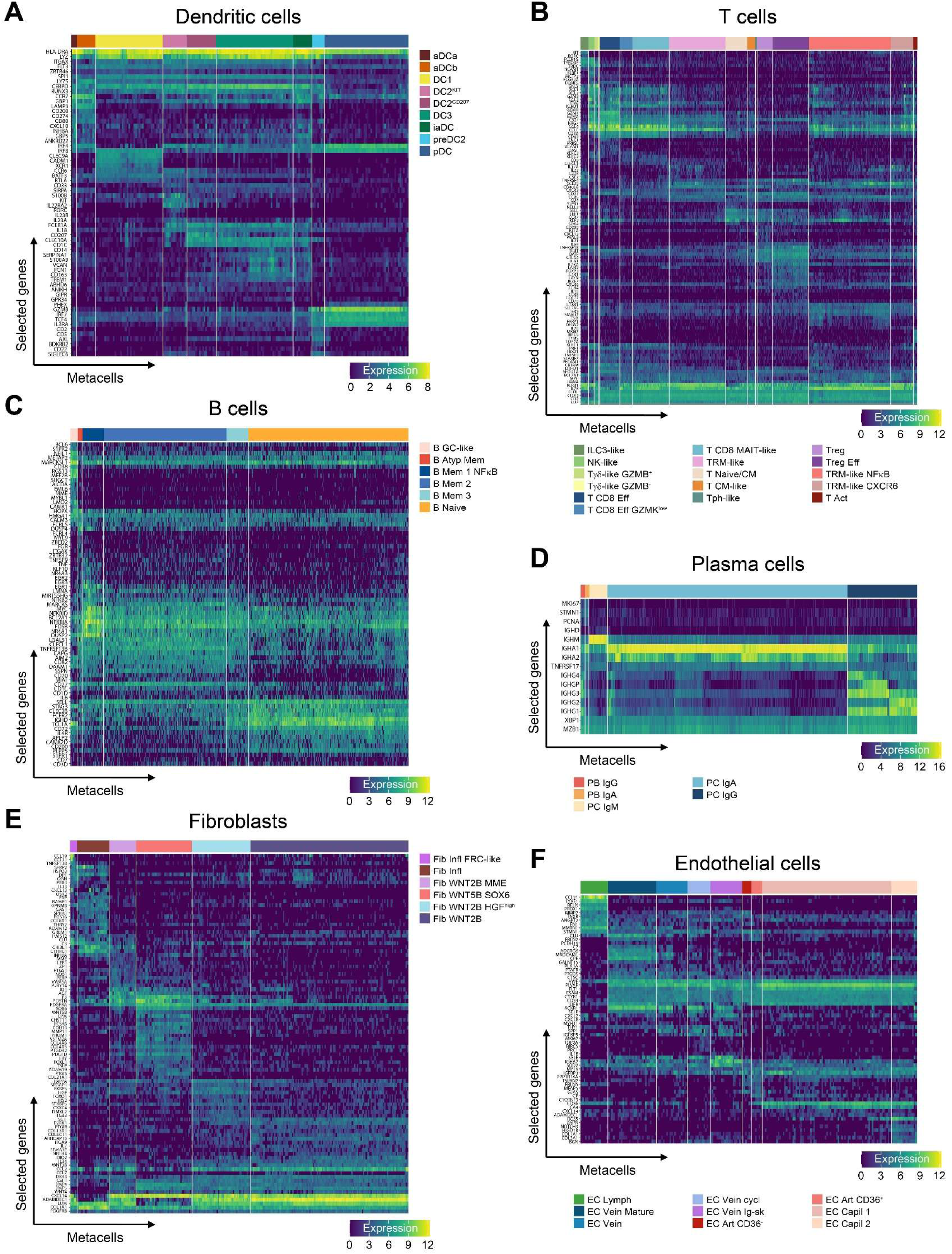
High-resolution analysis of cell lineages in CD inflamed ileums. **(A-F)** Heatmap showing expression of representative genes (rows) in single metacell (columns) grouped as indicated within dendritic cells (A), T cells (B), B cells (C), plasma cells (D), fibroblasts (E) and endothelial cells (F). A corresponds to the analysis of dendritic cells in MNP-enriched cell suspensions. (B-F) correspond to indicated lineage analyses in the CHU Nantes/Mount Sinai cohort.

**Figure S8:**
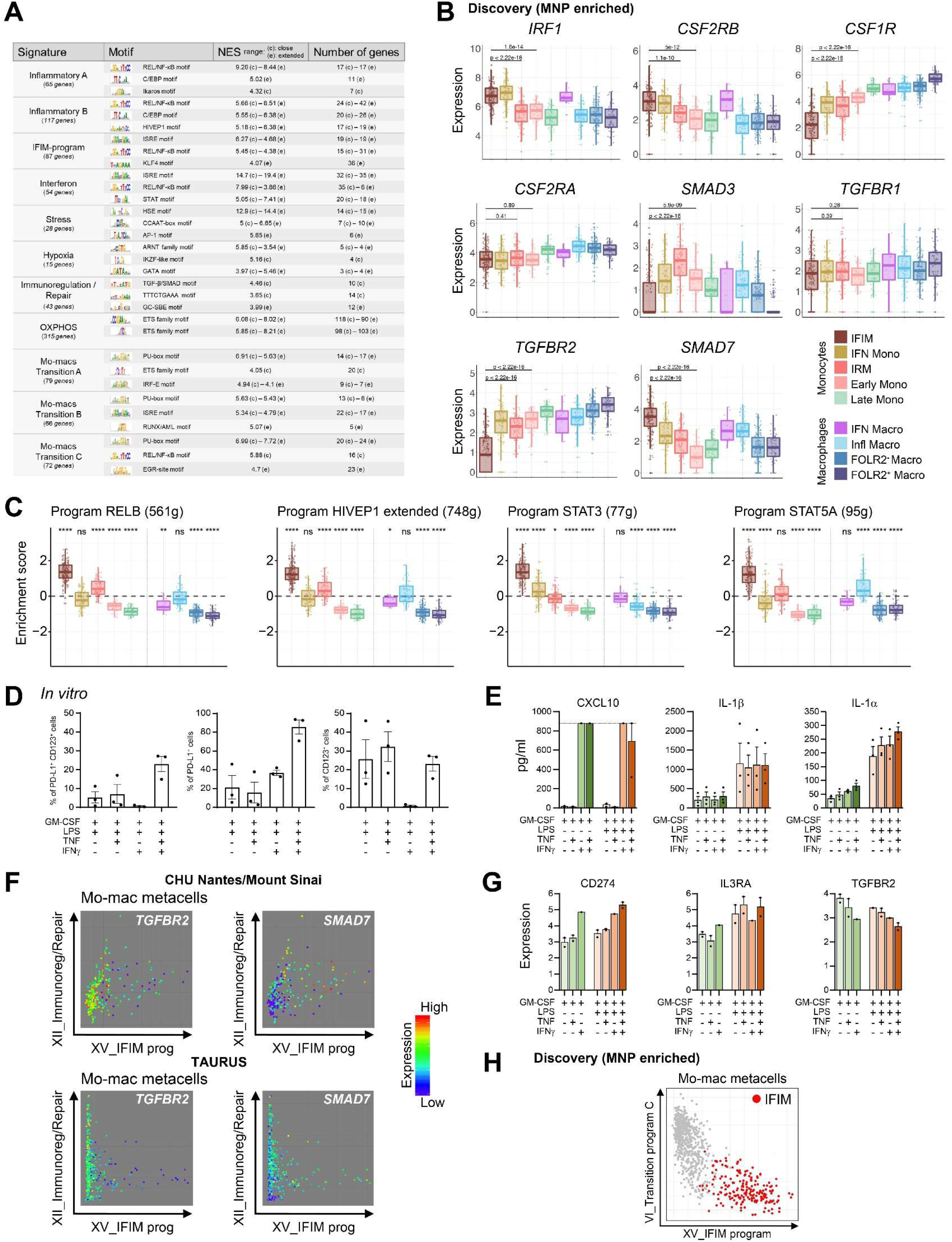
Molecular regulation of monocyte programs in CD ileums. **(A)** Top predicted TF motifs and Normalized Enrichment Score (NES) in the indicated mo-mac programs. **(B)** Boxplots showing expression levels of indicated genes across mo-mac groups. P-values were calculated using Mann-Whitney *U* tests with Bonferroni correction. For clarity, not all comparisons are shown. **(C)** Boxplots showing regulon activities of indicated TF calculated by SCENIC between mo-mac groups. P-values were calculated using Wilcoxon signed-rank tests (group median vs. global mean) with Bonferroni correction between all groups. *\*P < 0.05, **P < 0.01*, *\*\*\*P < 0.001, ****P < 0.0001, n.s. non-significant* (**Table S28**). **(D)** Bar plots (mean +/−s.e.m.) of monocyte phenotypes after culture in the indicated conditions. **(E)** Bar plots (mean +/−s.e.m.) showing the concentrations of the indicated cytokines in supernatants collected after 5 days of monocyte culture in the indicated conditions. **(F)** Color-coded expressions of *TGFBR2* and *SMAD7* in mo-mac metacells spread onto scatterplots according to their expression of programs “Immunoregulation/Repair” (y-axis) and “IFIM-program” (x-axis) in the CHU Nantes/Mount Sinai (left) and TAURUS (right) cohorts. **(G)** Box plots showing the expression of indicated genes in blood monocytes from healthy donors analyzed by RNAseq after 5 days of culture *in vitro* in the indicated conditions (n=3). **(H)** Scatterplots of total mo-mac metacells according to their expression of programs “Transition_C” (y-axis) and “IFIM-program” (x-axis).

**Figure S9:**
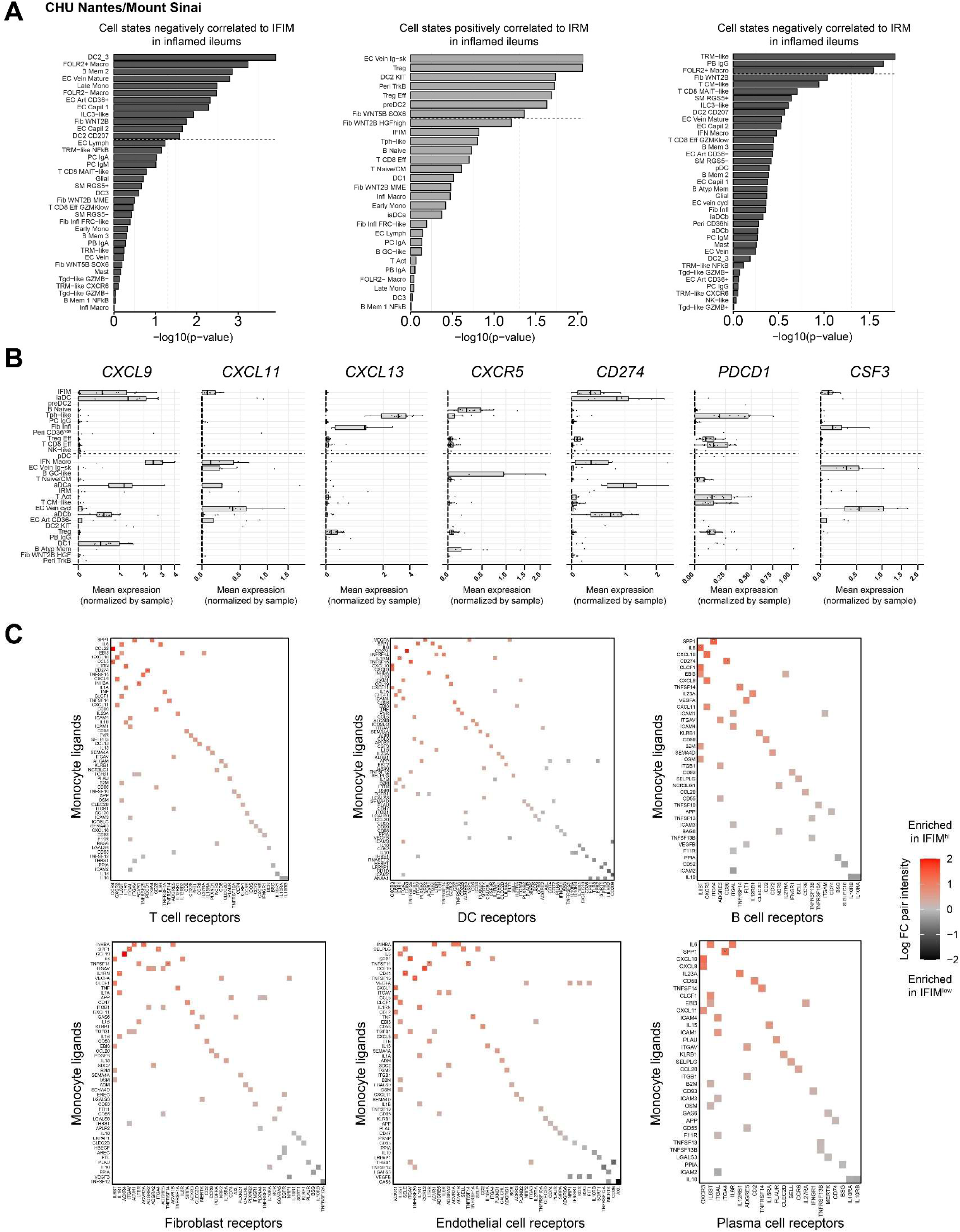
An IFIM-associated cytokine network at ulceration sites. **(A)** Bar plots showing scRNA-seq-defined cell states negatively correlated with IFIM (left), positively correlated to IRM (middle) or negatively correlated to IRM (right) in inflamed ileal tissue from Crohn’s disease (CD) patients. Cell states above the dashed line have a P-value < 0.05 (**Table S26**). **(B)** Expression levels of the indicated genes across cell states positively correlated with IFIM in inflamed CD ileums. **(C)** Comparative ligand-receptor network analysis between the two patient groups. Each square represents a validated pair of monocyte ligand (rows) to indicated cell type receptors (columns). The associated intensity (color-coded) is the log ratio of the difference in interaction score (the product between the mean expression of the ligand and the mean expression of the receptor) between IFIM^hi^ and IFIM^low^ samples. * denote pairs with a Benjamini-Hochberg adjusted P-value < 0.01 as estimated by permutation test.

**Figure S10:**
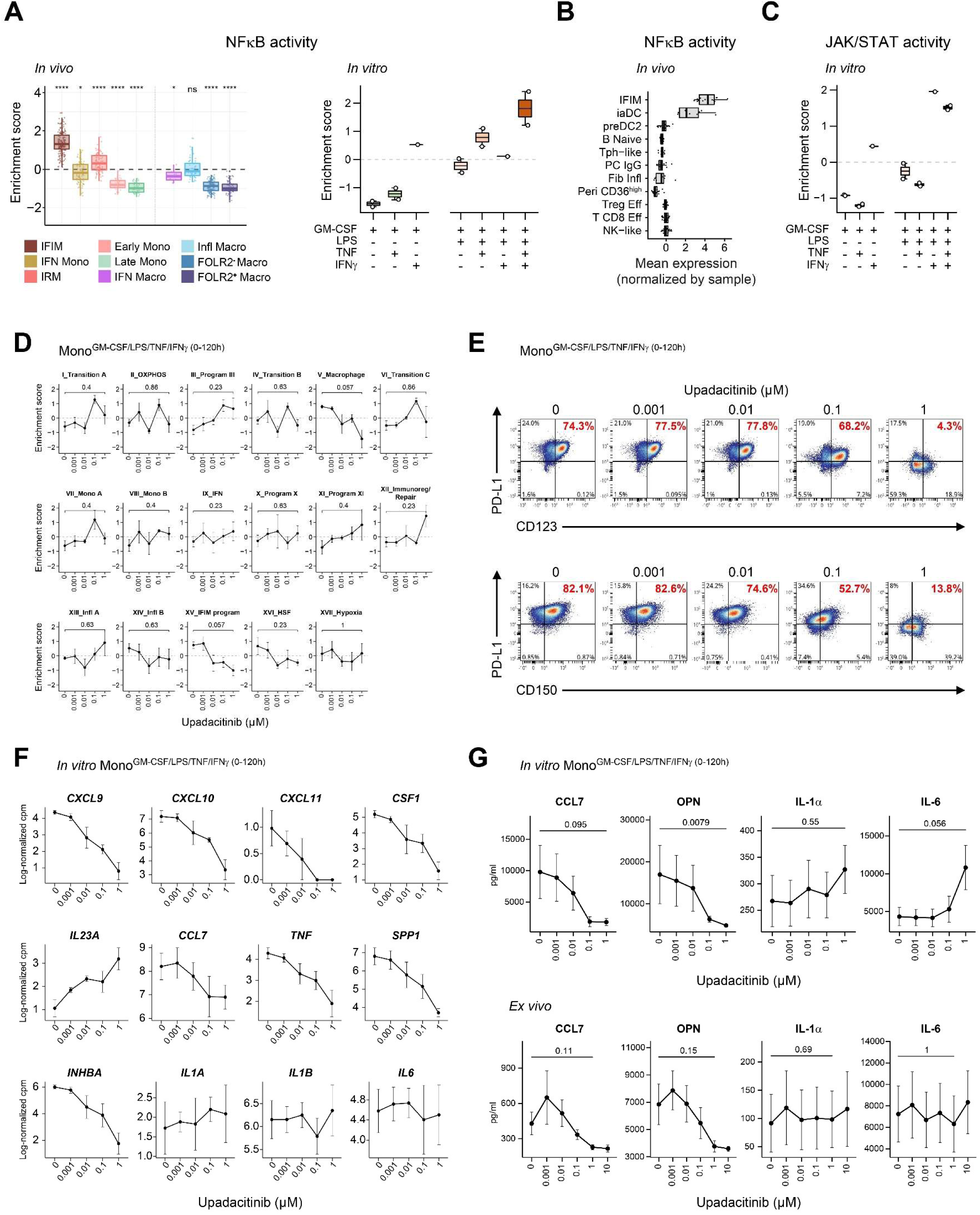
High JAK-STAT activation makes IFIM targetable by JAK inhibitors. **(A)** Boxplots comparing expression levels of predicted NFκB activity across mo-macs groups *in vivo* (left) and *in vitro* mono^GM-CSF/LPS/IFNγ/TNF(0-120h)^ (right). P-values were calculated using Wilcoxon signed-rank tests (group median vs. global mean) with Bonferroni correction between all groups. *\*P < 0.05, **P < 0.01, ***P < 0.001, ****P < 0.0001, n.s. non-significant* (**Table S30**). **(B)** Expression levels of the predicted NFκB activity across cell states positively correlated with IFIM in inflamed CD ileums. **(C)** Boxplots comparing expression levels of predicted JAK/STAT activity across mo-macs groups *in vitro* mono^GM-CSF/LPS/IFNγ/TNF(0-120h)^. **(D-E)** Mono^GM-CSF/LPS/IFNγ/TNF(0-120h)^ from healthy donors were exposed to upadacitinib at the indicated concentrations. (D) Dose effect of upadacitinib on the expressions of indicated scRNA-seq-defined mo-mac programs assessed by RNAseq. (E) Representative flow cytometry dot plots of dose effect of upadacitinib on monocytes phenotypes. **(F)** Dose effect of Upadacitinib on the expression of indicated genes assessed by RNAseq. **(G)** Dose effect of upadacitinib on cytokines levels detected in supernatants of (top) monocytes differentiated as in (D-E), (bottom) *lamina propria* cells isolated from surgically resected inflamed ileums of IFIM^high^ CD patients after 2 days of culture.

## Notes

### Competing Interest Statement

The authors have declared no competing interest.

